# Beyond pathologist-level annotation of large-scale cancer histology for semantic segmentation using immunofluorescence restaining

**DOI:** 10.1101/2022.05.09.489968

**Authors:** Daisuke Komura, Takumi Onoyama, Koki Shinbo, Hiroto Odaka, Minako Hayakawa, Mieko Ochi, Ranny Herdiantoputri, Kei Sakamoto, Hiroto Katoh, Tohru Ikeda, Tetsuo Ushiku, Shumpei Ishikawa

## Abstract

Numerous cancer histopathology specimens have been collected and digitised as whole slide images over the past few decades. A comprehensive evaluation of the distribution of various cells in a section of tumour tissue can provide valuable information for understanding cancer and making accurate cancer diagnoses. Deep learning is one of the most suitable techniques to achieve these goals; however, the collection of large, unbiased training data has been a barrier to producing accurate segmentation models. Here, we developed a pipeline to generate SegPath, the largest annotation dataset that is over one order of magnitude larger than publicly available annotations, for the segmentation of haematoxylin and eosin (H&E)-stained sections for eight major cell types. The pipeline used H&E-stained sections that were destained and subsequently immunofluorescence-stained with carefully selected antibodies. The results showed that SegPath is comparable to, or significantly outperforms, conventional pathologist annotations. Moreover, we revealed that annotations by pathologists are biased toward typical morphologies; however, the model trained on SegPath can overcome this limitation. Our results provide foundational datasets for the histopathology machine learning community.

---

Tumour tissues comprise various cell types, each with a unique function and morphology. In cancer histopathology, information on cell components and their distribution in the tumour tissues of patients helps with diagnoses, the classification of tumour subtypes, prediction of prognosis and therapeutic effects, and understanding the underlying mechanisms of carcinogenesis ^1, 2^. Although pathologists approximately estimate such information in clinical practice, it is almost impossible to measure cell components and distribution data quantitatively and comprehensively, particularly for large tissue specimens. Therefore, an automatic segmentation system using routinely used haematoxylin and eosin (H&E)-stained tumour sections can be highly valuable in medical practice and cancer research.

Deep neural networks are emerging machine-learning technologies capable of performing such tasks with remarkable accuracy^3–6^. However, the high performance of deep neural networks is because of their abundant annotations, which are often difficult to obtain in medical imaging. For the semantic segmentation of H&E images, there have been many efforts to annotate tissues or cells, most of which rely on human annotators^6, 7^. However, manually annotating tumour tissues is not feasible by non-pathologists and requires considerable time and labour, limiting the generation of large-scale annotated datasets. Another important issue that has often been overlooked in previous research is that human annotations may not cover the full diversity of cell morphologies. Cells do not always have the typical morphologies described in textbooks. The surrounding environment (e.g., narrow lumen) can deform cells, which may present atypical morphologies depending on the location and angle of the cell cross-section. Cells can also change their morphologies due to molecular interactions with the surrounding microenvironment. For example, identifying tumour vascular endothelial cells may be difficult given their enlarged nuclei and morphologies, which are similar to those of epithelial cells^8^. Additionally, some cell types, such as myeloid cells, can be extremely difficult for pathologists to accurately identify, as evidenced by the high rate of macrophage count discordance among pathologists^9^. These factors prevent the accurate annotation of cells with atypical morphologies by pathologists and potentially limit the performance of segmentation models trained on the datasets.

Recently, new methods utilising special stains for the annotation of histological images have emerged to overcome this limitation. Haan *et al*.^10^ developed a deep learning-based method to transform renal H&E-stained sections into special- stained sections, such as Masson’s trichrome and periodic acid–Schiff, to help pathologists with making diagnoses. By combining two networks, namely, a generator network to virtually stain unlabelled auto-fluorescent kidney tissues with H&E or special stains and stain transformation network to transform H&E images into special stain images, the method does not need to rely on unpaired images, which can degrade the transformation performance. This approach is useful for virtual staining, but its utility for cell-type segmentation remains unclear. Additionally, stain transformation uses virtually-stained rather than real H&E images in principle; thus, it can introduce prediction errors derived from the virtual stain generator network, although it works in renal pathology. Bulten *et al*.^11^ created a dataset for epithelium segmentation of H&E-stained prostate specimens using immunohistochemistry-restained images as a guide for generating masks. This approach is powerful as the H&E images are real, and the restaining procedure can produce perfectly-matched slides instead of consecutive slides, enabling accurate segmentation masks without the need for pathologists. The limitation of their study, however, was that they applied this method to only the epithelia of prostate tissues.

Our study has extended the aforementioned approach by creating a dataset for the semantic segmentation of various tissues or cells at an unprecedented scale. We developed an annotation workflow with minimal pathologist intervention based on H&E-stained sections that were destained and immunofluorescence (IF)-stained. Because IF relies on the proteins expressed in target cells, it can capture the target cells with diverse morphologies in a better manner than human annotations. Using carefully selected antibodies with high specificities for each of the eight major constituent cells in tumour tissues, we generated SegPath, a high-quality dataset of diverse cell types. SegPath is the largest annotated tissue and cell segmentation dataset with H&E images of various organs. SegPath will be made public to contribute to the development of new segmentation models using this dataset.

## RESULTS

### Dataset generation workflow

The workflow for creating SegPath is shown in Fig. 1a. First, tissue microarray (TMA) sections were stained with H&E using a standard procedure. They were then digitised using a slide scanner to create whole slide images (WSIs). After destaining the H&E-stained sections with alcohol and autoclave processing, IF and 4’,6-diamidino-2- phenylindole dihydrochloride (DAPI) nuclear staining were performed using antibodies that specifically recognised each cell type. The sections were then digitised again. Multiresolution rigid registration between the H&E and IF images was performed to ensure that the haematoxylin component in the H&E images and DAPI in the IF images, both recognising nuclei, had been aligned. Registration was first performed at the WSI level and then at the patch level. After rigid registration, we observed that a few cells shifted slightly in inconsistent directions during destaining and restaining. Therefore, we performed an additional nonrigid registration for each patch to accurately align the nuclei (Supplementary Fig. 1a).

**Fig. 1.**
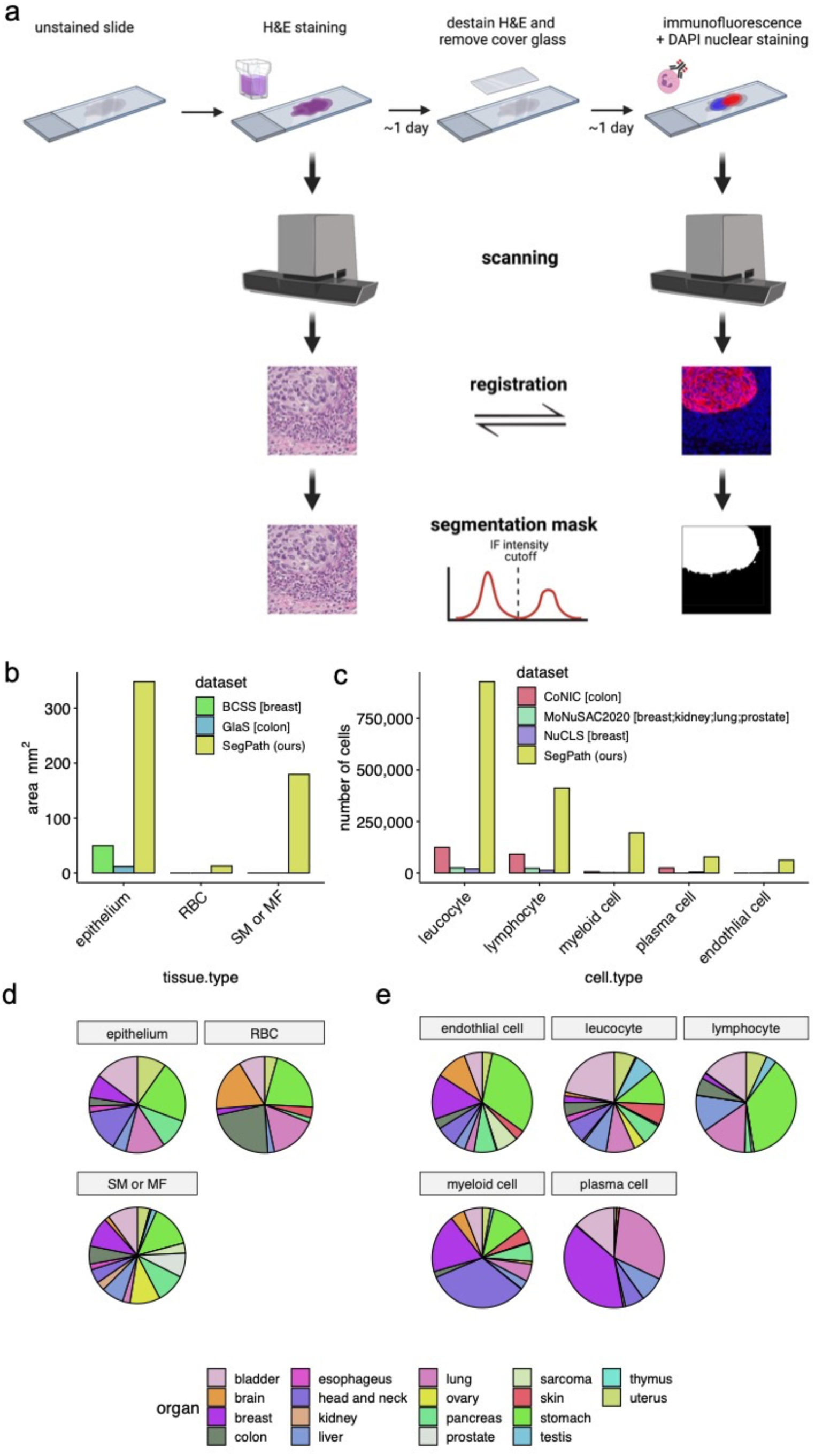
Generation of annotation masks for tissue/cell-type segmentation using IF restaining. **a,** Workflow overview. After scanning the H&E-stained sections, the sections were destained and restained with DAPI nuclear staining and IF staining with target-specific antibodies. The slides were then scanned, and the positions of the two slides were aligned with registration algorithms. Small patches were created. Cut-off values of IF signal intensity were determined for each patch to generate segmentation masks in an iterative manner based on the segmentation results of the deep neural network training on the generated masks. For endothelial cells, leucocytes, lymphocytes, myeloid cells, and plasma cells, a nucleus detection algorithm was applied to the DAPI channel. Positive signals of the target cell in IF were transferred to the corresponding nuclei. **b,** Annotated areas for each tissue and **c,** the number of annotated cells for each cell type in SegPath. Those in publicly available datasets (BCSS^13^, GlaS^15^, CoNIC^5^, MoNuSAC2020^14^, and NuCLS^7^) are also shown. Organs in brackets represent the target organs of the dataset. ‘SM’ or ‘MF’ includes all stroma in the GlaS and BCSS datasets. **d,** Distribution of target organs in SegPath. **e,** Distribution of cell types in SegPath. SM, smooth muscle cell; MF, myofibroblast.

Next, we created the binary segmentation mask based on the IF images (hereafter referred to as ‘IF-mask’). The area in which the fluorescence intensity exceeded the cut-off value was labelled as positive. To remove the false positives from red blood cell (RBC) autofluorescence, positive regions estimated using the deep neural network, which was trained on the dataset using an anti-CD235a antibody recognising RBCs, were removed from the non-RBC datasets. For the haematopoietic and endothelial cells, the segmentation masks of the target cells were transferred to the target cell nuclei to make the segmentation task more tractable. We used Cellpose^12^, a deep neural network model for nuclear segmentation, with the DAPI images to recognise the nuclei (Supplementary Fig. 1b). We then labelled whole nuclei as positive if the positive region overlapped with the nuclei over a certain level. Finally, we improved the fluorescence intensity threshold as a gradient was observed in the same WSIs; thus, the fixed threshold had not been optimal. The deep neural network model was trained on the dataset, and Otsu’s threshold for successfully segmented patches with positive regions was used in the next phase. For the other patches, the threshold was propagated from the neighbouring, successfully segmented patches. This process was repeated twice until the generated IF-masks had been converged (Supplementary Fig. 1c).

Nine different antibodies were used to cover the main cell components of the tumour tissue (Table 1). A mixture of anti-CD3 and anti-CD20 antibodies was used for lymphocytes. Because our workflow can generate segmentation masks in a high-throughput manner without the need for manual annotation, the size of the dataset is over one order of magnitude larger than the currently available segmentation mask datasets for tumour tissues^5, 7, 13–15^ (Fig. 1b, c). In addition, we created datasets for as many as 18 different organs from 1,583 patients (Fig. 1d, e) to cover a wider spectrum of cancer types (Supplementary Table 1) than in the currently available datasets, which cover up to four organs.

**Table 1.**
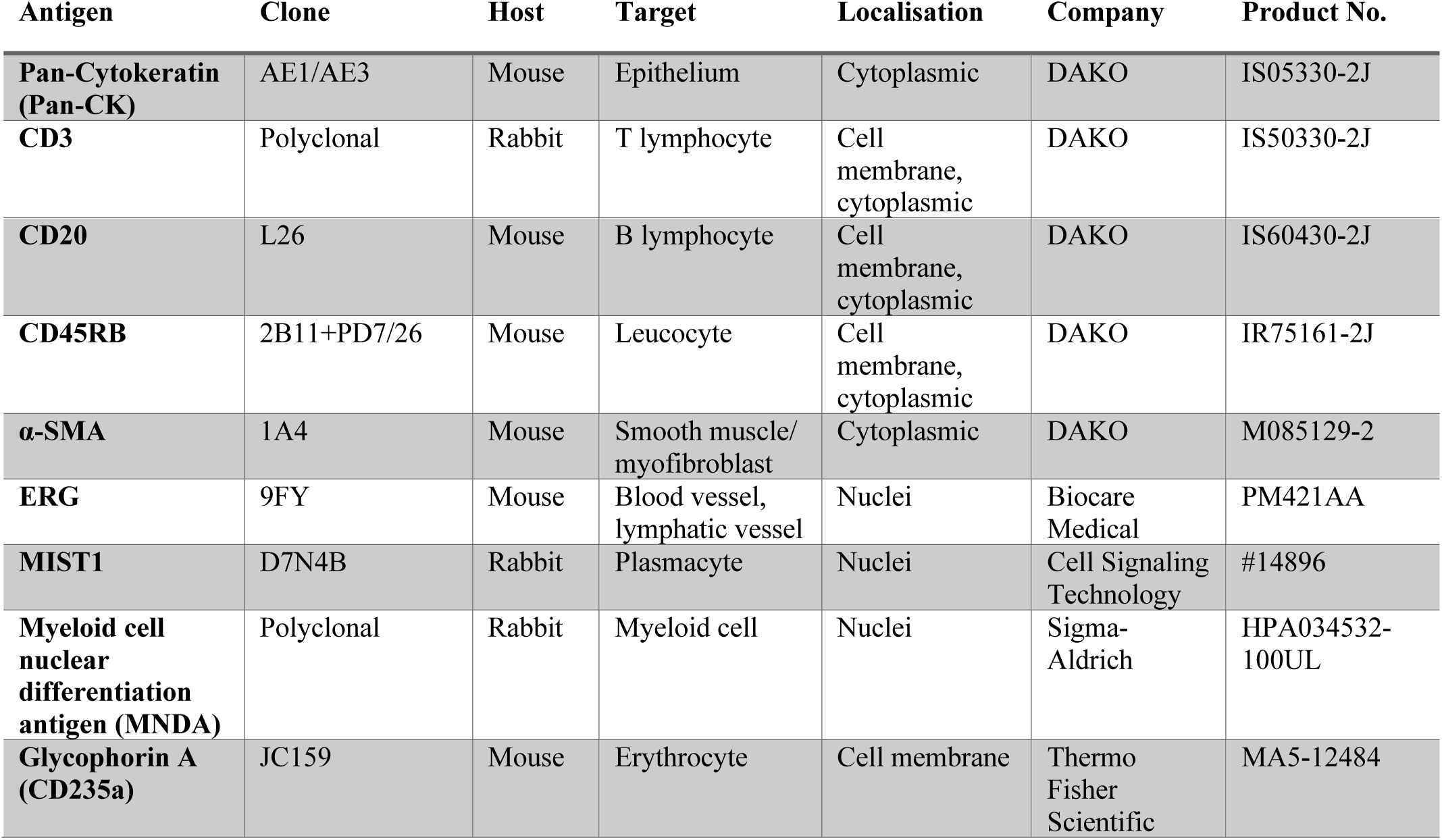
Antibodies used in this study.

### Antibody and organ selection

The choice of antibodies is one of the most important factors in the successful generation of IF-masks. We carefully selected the proteins that are specifically expressed in the target cell types based on the gene expression profiles (Fig. 2a). Moreover, we selected cytoplasmic proteins for tissue-type segmentation (epithelium and smooth muscle cell/myofibroblasts). For haematopoietic cells or the endothelia, we prioritised proteins localised in the nuclei of target cell types as the position of the cells can be easier to identify with the antibody to such proteins. For lymphocytes and leucocytes, we selected antibodies used in clinical practice that stained the cell membrane, mainly because the appropriate antibodies that localised to the nuclei could not be found despite various trials using candidate antibodies (data not shown).

**Fig. 2.**
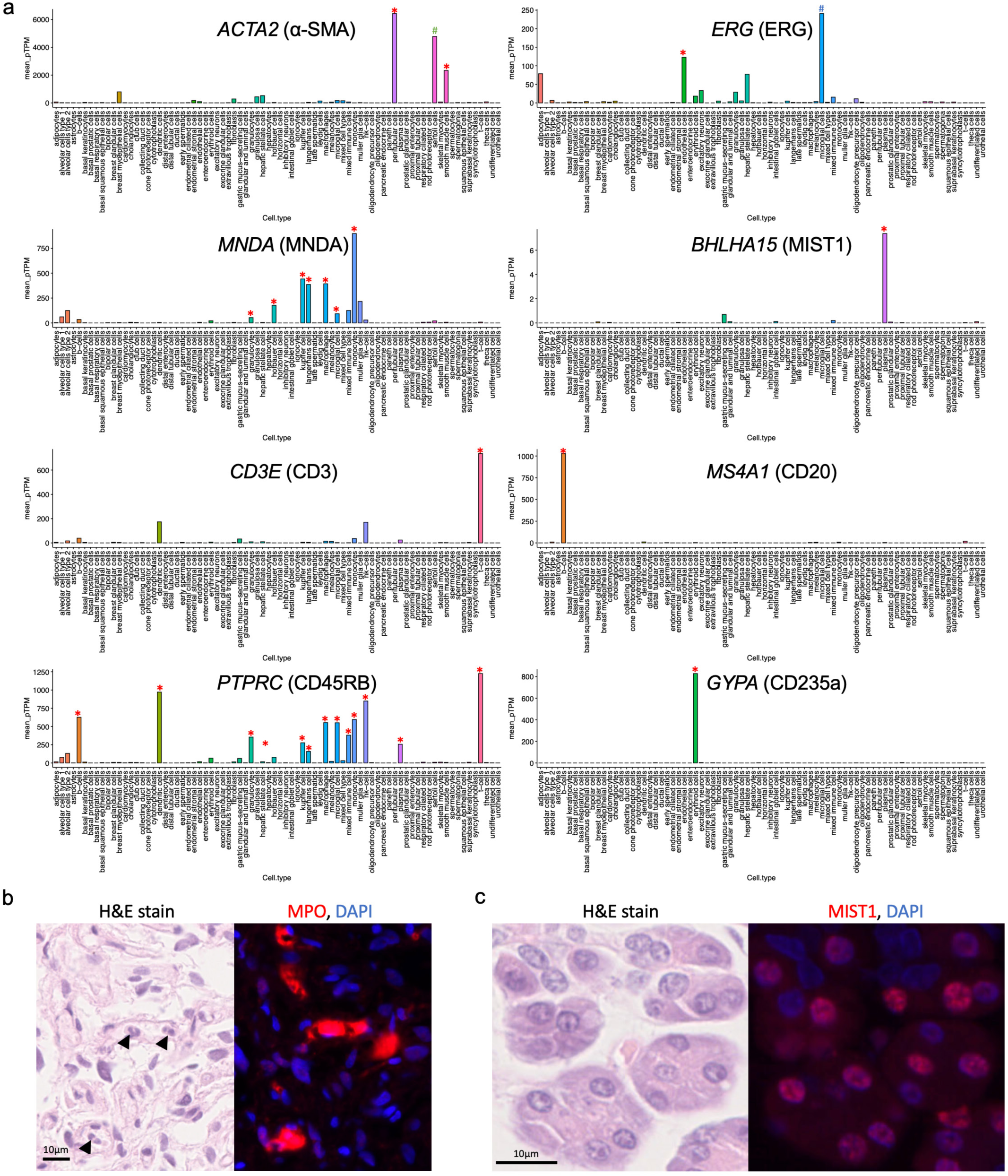
Selection of the antibodies and target tissues in SegPath. **a,** Gene expression specificities of selected antibodies. Gene expression data were retrieved from single-cell transcriptome profiles in the Human Protein Atlas^16, 17^. Target cell type is indicated by a red asterisk on the bar. *ACTA2* expression in Sertoli cells, indicated by a green octothorpe, was high in this dataset, but a pathologist could not confirm the positive staining of anti-α-smooth muscle actin (SMA) antibody; thus, testicular tissues were included in the dataset. *ERG* expression in microglial cells, indicated by a blue octothorpe, was higher in this dataset. This is highly likely to be an erroneous annotation of the single-cell transcriptome profile, as confirmed by a pathologist; thus, brain tissues were included in the dataset. **b,** H&E-stained image and the IF staining of anti-MPO antibody, which targets neutrophils. Antigens spread around the target cells, as indicated by arrowheads, prevent accurate mask generation. **c,** IF staining of anti-MIST1 antibody, which targets plasma cells. It unexpectedly stained the nuclei of some glandular epithelia, including the salivary gland and gastric epithelium. These tissues were excluded from SegPath.

We observed several failures during the antibody selection process. A few of the antibodies that had been initially selected exhibited low staining intensities or specificities, depending on the clone. In other cases, such as that with myeloperoxidase (MPO), the antigen diffused into the surrounding area, making it difficult to accurately identify the locations of cells (Fig. 2b). Although MIST1 is a plasmacytoid-specific antigen, it was slightly stained with the selected anti-MIST1 antibody in some glandular epithelial cells (Fig. 2c). Therefore, organs such as the stomach, pancreas, and salivary glands were excluded from the MIST1 dataset. After optimising the antibodies, the pathologists confirmed the specificity of IF staining of the target cells. Prostate and renal cancer and hepatocellular carcinoma cases were also removed from the pan-CK dataset due to the weak IF staining of tumour cells.

### Dataset evaluation

Figure 3 shows examples of the H&E-stained images, matched IF images, and generated IF-masks in SegPath for the selected antibodies in various organs. We verified that the target tissues or cells were specifically stained. For example, the anti-pan-CK antibody stained cytokeratin, which was localised in the cytoplasm of the epithelial cells. Although the nuclei of the epithelial cells were unstained in some cells, the borders of the epithelial tissue regions were clear, indicating the use of the mask for epithelial segmentation. The anti-αSMA antibody stained perivascular smooth muscle cells densely and smooth muscle or myofibroblasts in some stroma less densely (Supplementary Fig. 2), which possibly reflected the density and expression of αSMA (e.g., cancer-associated fibroblasts (CAFs) differentiate into cells with the myofibroblast phenotype, which are morphologically similar to smooth muscle cells but have variable αSMA expression depending on the degree of differentiation). Anti-CD45RB and anti-CD3/CD20 antibodies recognised the proteins on the cell membranes of leucocytes and lymphocytes, respectively; however, additional pre-processing using a nucleus detection algorithm made the generated masks cover the nuclei only, making the cell positions clearer. The masks were almost identical to the IF images for the antibodies against ERG, MNDA, and MIST1.

**Fig. 3.**
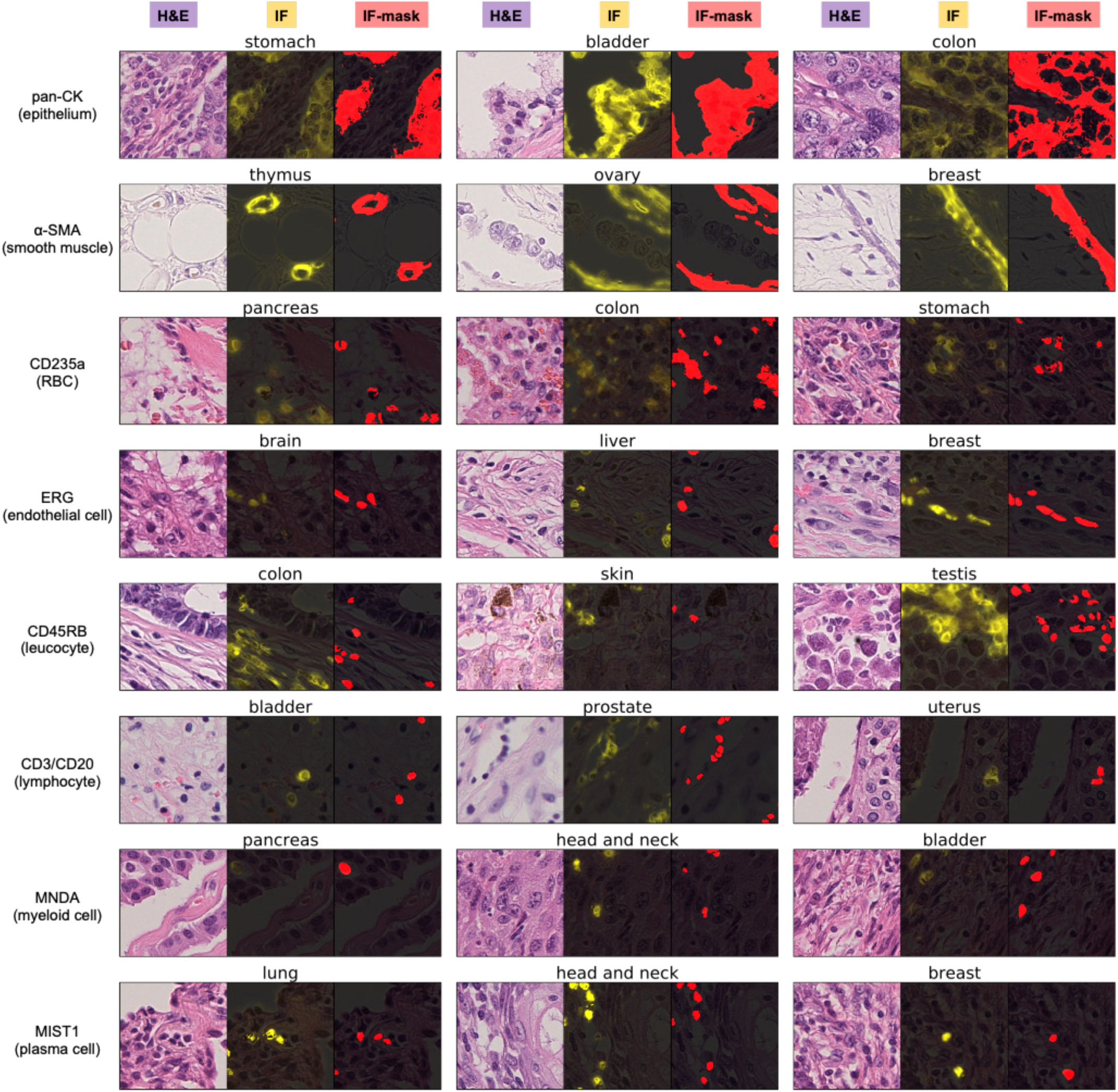
Generated masks in cancers of various organs. Each triplet shows an H&E-stained image, the corresponding registered IF image, and generated mask image (positive regions are indicated by red), from left to right, respectively. The organs are shown above each triplet. All image patches are 72.5 × 72.5 µm.

To quantitatively evaluate the quality of the IF-masks in SegPath, we compared them with two types of manual annotations: the annotation created by three trained pathologists evaluating the H&E images alone (hereafter referred to as ‘HE-path’), and both the H&E and corresponding IF images (pathologist-guided ground truth, hereafter referred to as ‘pGT’) (Fig. 4 and Extended Data Fig. 1). The evaluation dataset consisted of 10 image patches of 217.5 × 217.5 µm for each antibody. The pathologists generated HE-paths based on morphology and pGTs based on morphology and IF intensity and distribution. The regions or cells annotated by at least two pathologists were regarded as positive. Therefore, the HE-paths may be considered baselines in conventional manual annotations, and the pGTs can be considered closest to the ground truth as pathologists are thought to be less affected by spurious IF signals.

**Fig. 4.**
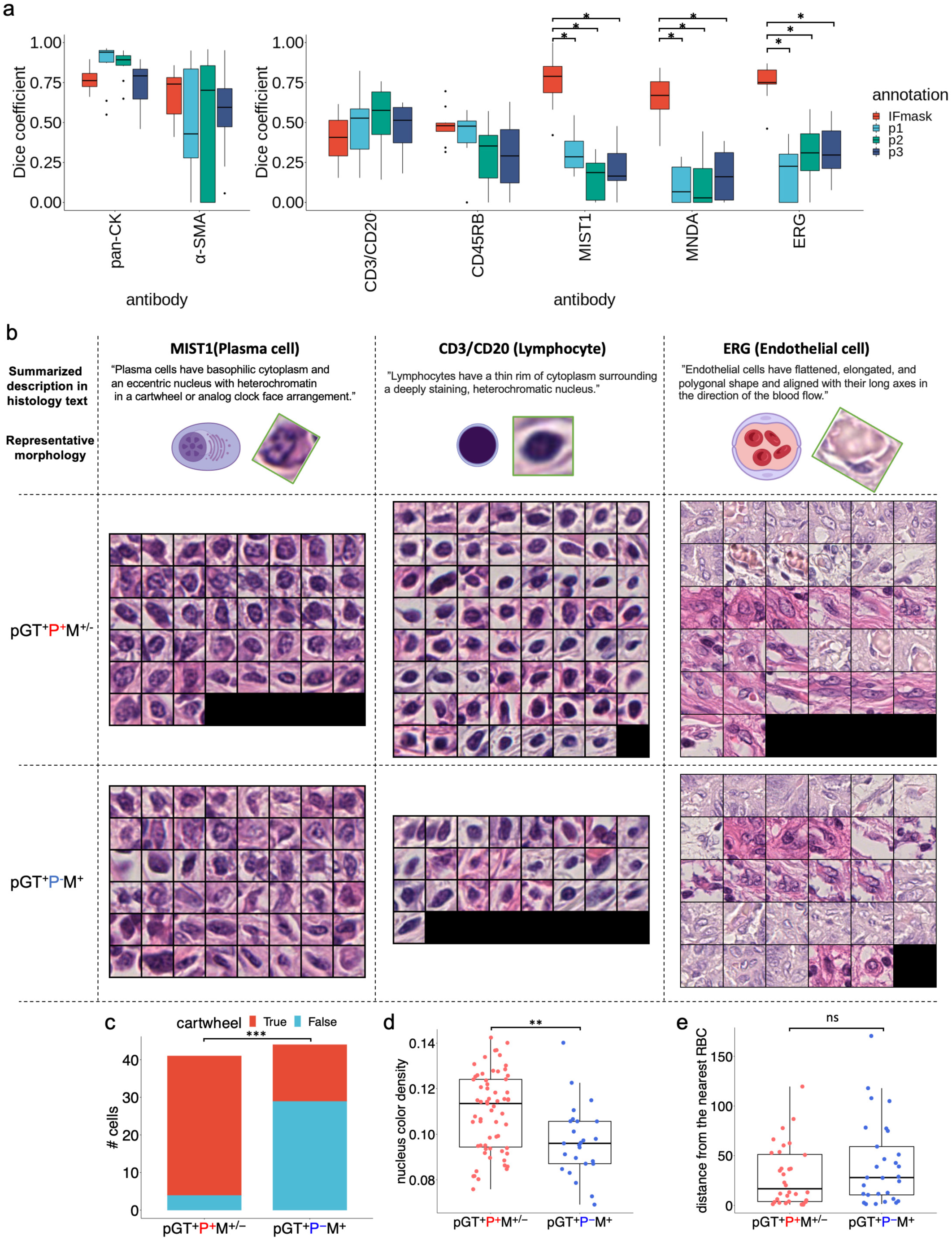
Evaluation of the annotation accuracy of SegPath. **a,** Annotation accuracy of pathologists and the generated masks in SegPath (*n* = 10 patches of 217.5 × 217.5 µm for each tissue or cell type) in comparison with pGT as ground truth. Dice coefficients of the IF-masks were compared with those of annotations by each pathologist. Two-sided Wilcoxon signed-rank test was used, and *P*-values were adjusted using the Benjamini–Hochberg method. *P* < 0.05 was considered statistically significant, as shown by asterisks. **b,** Ground truth (pGT) cell images annotated by multiple pathologists (pGT^+^P^+^M^+-^), and not identified by multiple pathologists but successfully annotated by the masks (pGT^+^P^-^ M^+^). The illustrations and the actual images of the representative cell morphologies and sentences describing the morphologies written in a histology textbook are shown in each cell type^18^. Original illustrations from BioRender were used except for the lymphocyte, whose nucleus was denser and larger than the original illustration. The image was adjusted to be more similar to the representative morphology. For the box plot, the lower and upper hinges correspond to the 25^th^ and 75^th^ percentiles, respectively; the upper whisker extends from the hinge to the largest value no further than 1.5 × IQR from the hinge. The lower whisker extends from the hinge to the smallest value at 1.5 × IQR of the hinge. pGT, ground truth; P, HE-path; M, IF-mask. **c,** Distribution of plasma cells with or without the typical cartwheel-shaped nuclei (*n* = 41 cells for pGT^+^P^+^M^+-^ and *n* = 44 cells for pGT^+^P^-^M^+^, two-sided Fisher’s exact test). **d,** Nucleus haematoxylin intensity of lymphocytes (*n* = 63 cells for pGT^+^P^+^M^+-^ and *n* = 25 cells for pGT^+^P^-^M^+^, two-sided Mann– Whitney *U*-test). **e,** Distance (µm) from the endothelial cell to the closest RBC (*n* = 32 cells for pGT^+^P^+^M^+-^ and *n* = 29 cells for pGT^+^P^-^M^+^). *\*\*\*P<*0.0001, *\*\*P<*0.01

First, we examined the concordance of the HE-paths among the three pathologists (Extended Data Fig. 2a). The concordance varied immensely depending on the tissue and cell type. It was nearly identical in epithelial tissues but showed little overlap in the endothelia, plasma cells, and myeloid cells. This highlighted the difficulty faced by pathologists when accurately identifying these cells.

Next, we evaluated the correctness of the HE-paths and IF-masks of each tissue or cell type in terms of the Dice (F1 score), precision, and recall (Fig. 4a and Extended Data Fig. 2b, c) indices compared to those of the pGTs. We found that the performance of the HE-paths for the five cell types was low, indicating that it would be difficult for pathologists to identify such cells accurately. By contrast, the IF-masks were substantially more accurate than the HE- paths, especially in plasma (MIST1), myeloid (MNDA), and endothelial (ERG) cells. Unlike the HE-paths, the performance of leucocytes (CD45RB) and lymphocytes (CD3/CD20) was lower than that of the other three cell types. This may have been because of the antibodies used to recognise the proteins in the cell membrane. This made it difficult to estimate the exact locations of the cells, especially with the variable intensity of the staining. Nevertheless, the performance of leucocytes and lymphocytes was comparable to that of the HE-paths.

We hypothesised that pathologists could not accurately identify cells with atypical morphologies. To clarify the biases in the annotations of the pathologists, we analysed images of cells that the pathologists correctly identified (pGT^+^P^+^M^+-^) and those that the pathologists could not correctly identify, but that the IF-masks could identify (pGT^+^P^-^ M^+^) (Fig. 4b and Extended Data Fig. 3). Although some of the cells missed by the pathologists may have been missed because they were overlooked, the morphological characteristics of the cells that the pathologists correctly identified were clarified. Overall, the shapes and sizes of pGT^+^P^+^M^+-^ cells were more uniform than those of pGT^+^P^-^M^+^ cells, implying a bias in the decisions of the pathologists toward the typical morphologies. Furthermore, we quantitatively investigated how pGT^+^P^+^M^+-^ morphology is biased toward textbook descriptions. For example, plasma cells are generally described as having a basophilic cytoplasm and an eccentric nucleus with heterochromatin in a characteristic cartwheel or clock face arrangement. As expected, the plasma cells in pGT^+^P^+^M^+-^ tended to have cartwheel-shaped nuclei (Fig. 4c), but less so in pGT^+^P^-^M^+^ cells, suggesting that pathologists cannot accurately identify plasma cells without clear cartwheel-shaped nuclei. By contrast, the basophilicity of the cytoplasm and eccentricity of the nucleus were not significantly different between pGT^+^P^+^M^+-^ and pGT^+^P^-^M^+^ cells (data not shown). Lymphocytes are generally characterised by a high nuclear–cytoplasmic ratio and dense nuclei. However, the lymphocytes overlooked by the pathologists often had thinner nuclei (Fig. 4d and Supplementary Fig. 3). There were no significant differences in the shapes of the vascular endothelial cells, but they were more likely to be correctly recognised if they were surrounded by multiple RBCs (Fig. 4e). With the myeloid cells, the pathologists were unlikely to miss polymorphonuclear leucocytes, such as neutrophils, as they are easy to identify (Extended Data Fig. 3).

The morphologies of cells that presented false negatives in the IF-masks but true positives in the HE-paths (pGT^+^P^+^M^-^) were also examined (Extended Data Fig. 4). The results showed that most of the false negatives were due to the lack of false negatives for cell nuclei detection by Cellpose. The reason for this is unclear, but it may be due to the accuracy of the deep learning model used in Cellpose. However, morphological bias was unclear on visual inspection.

In summary, we revealed an inherent morphological bias in the annotations of the pathologists. However, the SegPath annotations are likely to be less prone to such bias and may enable the production of accurate segmentation models to cover the morphological diversity of cells.

### Segmentation model trained on the dataset

We generated many annotated histological images of various tissues or cell types with diverse morphologies. To investigate whether such large-scale datasets improve segmentation performance, we trained semantic segmentation models on the training set of SegPath for each cell type independently using a convolutional neural network. We then tested one part of the test dataset, which was the same as that used in the previous section. We evaluated the segmentation performance gains as a function of increasing patches for epithelia, smooth muscle cells/myofibroblasts, and the number of endothelial cells, leucocytes, lymphocytes, plasma cells, and myeloid cells (Fig. 5). Similar to other image classification tasks for pathological images^19^, the predictive performance increased as more samples were used for model training. Apart from RBCs (CD235a) and lymphocytes (CD3/CD20), the performance gain did not seem to be saturated, indicating that more annotations can improve the segmentation performance. This result indicates the importance of our approach in obtaining a large number of annotated images in a high-throughput manner with minimal pathologist intervention.

**Fig. 5.**
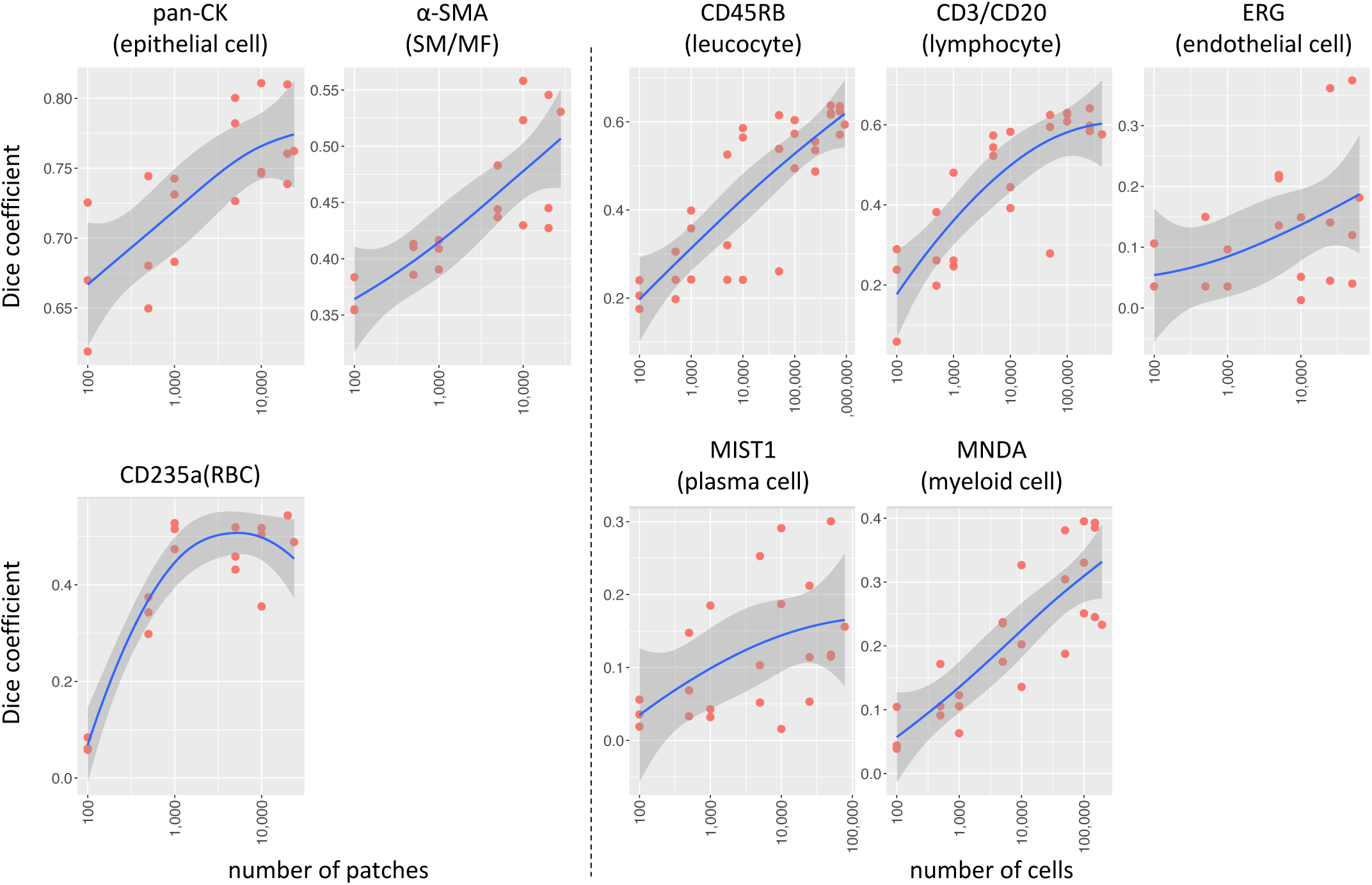
Effect of training sample sizes on segmentation models. Each point represents the Dice coefficient (F1 score) for each training on the same test dataset. SM, smooth muscle; MF, myofibroblast.

Next, we evaluated the performance of the segmentation models trained on the full training dataset (Fig. 6a) in SegPath and tested it on the same part of the test dataset, as described above. We observed that the overall performance of the segmentation models was comparable to that of the pathologists (HE-path) assessing the epithelium and leucocytes and, surprisingly, better than that of the pathologists assessing the other tissues or cell types in terms of the Dice coefficient. Cells that were not identified by the pathologists but identified by the trained models are shown in Extended Data Fig. 5. Similar to the IF-masks, plasma cells without typical cartwheel-shaped nuclei (Fig. 6b) and lymphocytes with thin nuclei were detected by the segmentation models more often than the HE-paths (Fig. 6c, Supplementary Fig. 4). These results indicate that the datasets enable the segmentation models to cover diverse morphologies. As shown in the epithelial cells in Fig. 6d, the segmentation models could identify even small areas that are difficult to identify. These results may be useful in cases of solitary cancer cells, such as those in diffuse-type gastric cancer (Supplementary Fig. 5). As shown in Fig. 6d, the segmentation models were able to identify smooth muscle around blood vessels, which is normally difficult to identify, possibly due to the lack of clear boundaries within the surrounding tissue.

**Fig. 6.**
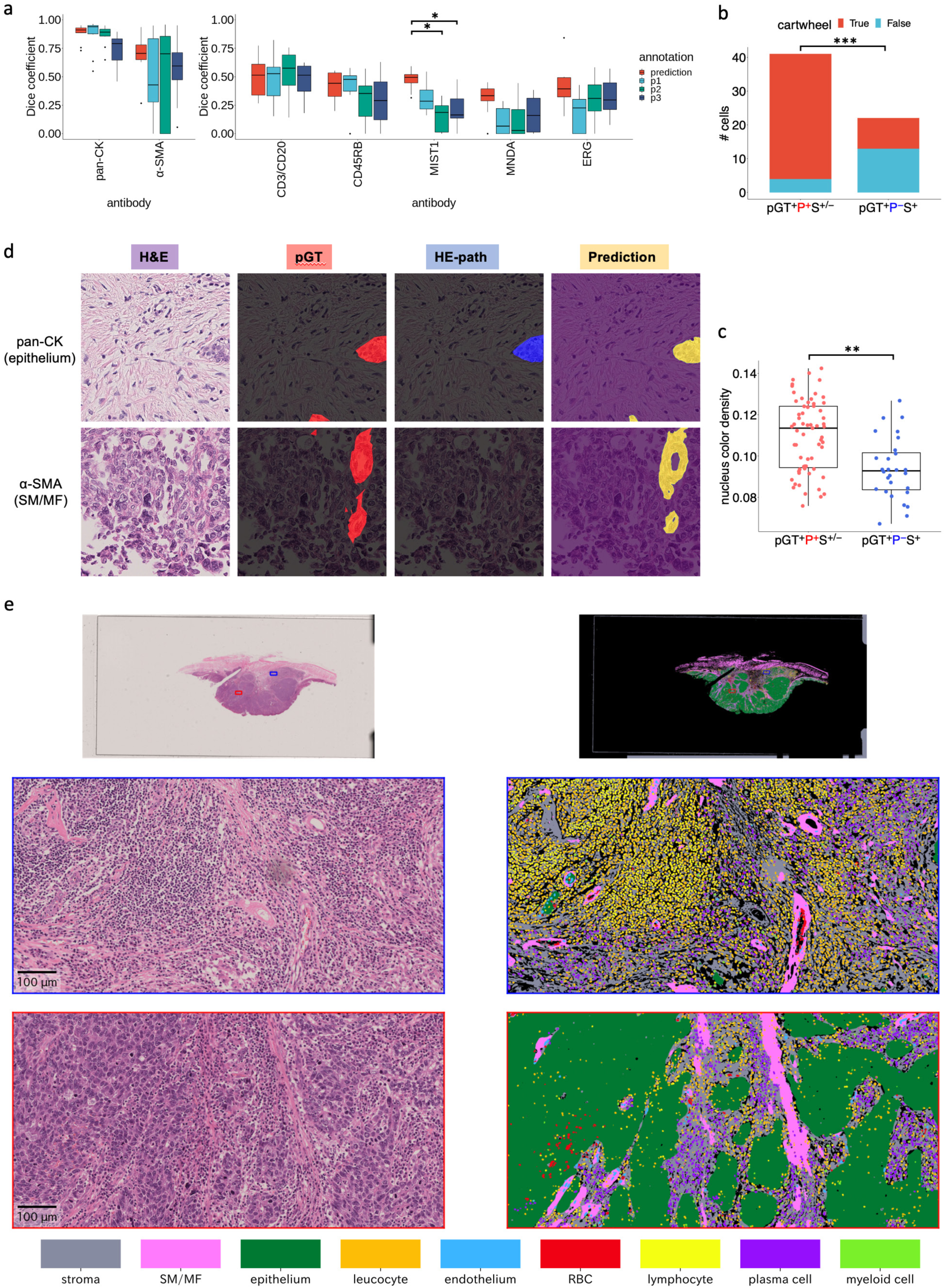
Performance evaluation of the segmentation models trained on the generated annotation masks. **a,** Comparison of the annotation accuracies between pathologists and prediction of the segmentation models in terms of the Dice coefficient (F1 score), (*n* = 10 patches of 217.5 × 217.5 µm for each tissue or cell type). pGT annotations were made by pathologists who evaluated both H&E and the corresponding IF images. Regions or cells annotated by at least 2 of the 3 pathologists were used. **b,** Distribution of plasma cells with or without typical cartwheel-shaped nuclei (*n* = 41 cells for pGT^+^P^+^S^+-^ and *n* = 22 cells for pGT^+^P^-^S^+^). **c,** Nuclear haematoxylin intensity of lymphocytes (*n* = 63 cells for pGT^+^P^+^S^+-^ and *n*=28 cells for pGT^+^P^-^S^+^). For the box plot, the lower and upper hinges correspond to the 25^th^ and 75^th^ percentiles, respectively; the upper whisker extends from the hinge to the largest value no further than 1.5 × IQR from the hinge. The lower whisker extends from the hinge to the smallest value at 1.5 × IQR of the hinge. pGT, ground truth; P, HE-path; S, Prediction by the segmentation model. **d,** Comparison of pathologist annotations for AE1/3 and SMA **e,** WSI level segmentation of a gastric cancer specimen using the segmentation models. Regions in blue and red rectangles in the top WSI level image are enlarged in the middle and bottom images with a bounding box of the same colour. Different colours represent different cell types as indicated. Grey regions annotated as stroma were tissue regions not detected by any segmentation models. SM, smooth muscle; MF, myofibroblast.

Finally, the segmentation models were applied to external gigapixel WSIs from various cancer tissues (Fig. 6e, Extended Data Fig. 6–8). To combine the outputs of the segmentation models for each tissue or cell type to generate a multi-tissue/cell segmentation result, we utilised the cell lineage hierarchy (see Methods for details), such that leucocytes included lymphocytes, myeloid cells, and plasma cells, and the tissue regions not positive by any models were labelled as ‘stroma’. Thus, we successfully generated segmentation results for nine tissues and cell types. The densities of the predicted smooth muscle/myofibroblast regions were variable; perivascular smooth muscle cells were dense, but other stromal regions were less dense, as shown in Extended Data Fig. 7b. This reflected the expression of αSMA in smooth muscle cells or CAFs with myofibroblast phenotypes as discussed above. The model was likely to capture the characteristic structures of various tumours, including benign and malignant salivary gland tumours, which were not included in the training data. For example, lymphoid structures filled with many lymphocytes, vascular wall linings with endothelial cells surrounded by smooth muscle cells and containing RBCs, and the rich infiltration of plasma cells around cancer cells were detected (Fig. 6e). Additionally, the infiltration of small cancer foci with no apparent glandular formation was successfully identified in the gastric cancer specimen (Extended Data Fig. 6a). A dense lymphoid stroma and double layer of oncocytic epithelia were also successfully identified in the Warthin’s tumour specimen (Extended Data Fig. 8a).

## DISCUSSION

Pathologists and biological researchers have routinely used immunohistochemistry to identify specific cells or tissues in research and clinical practice because of its simplicity and accuracy when appropriate antibodies are used. This study approached the problem of annotation generation for tissue or cell segmentation by leveraging immunostaining with cell-specific or tissue-specific antibodies. We generated SegPath, an accurate and high-volume dataset for the tissue or cell segmentation of H&E images based on a newly developed workflow that utilises IF staining. In SegPath, we targeted eight cell or tissue types that constitute the major component in tumour microenvironment^20^, and the granularity in the cell hierarchy was determined based on the potential feasibility of segmentation in H&E images. The advantages of our workflow were that identical sections were stained with H&E and IF, which enabled the precise localisation of target tissues or cell types. Furthermore, higher annotation accuracy could be achieved even if the target tissues or cells presented atypical morphologies. A series of experiments showed that the generated masks and segmentation models trained on the dataset achieved good performance with various morphologies. Although each image contained a mask for only one tissue or cell type, multiple cell types or tissues may be segmented using the outputs from multiple models, as shown in the last experiment.

Our experiment revealed that pathologists could miss or make incorrect labels with variable extents depending on the cell type. Furthermore, the annotations of the pathologists were biased toward typical morphologies. Cells with atypical morphologies and/or surrounding microenvironments may be subtypes with unique functions or states, as suggested by previous studies^21, 22^. The datasets of existing studies based on annotations by pathologists also contained biases toward typical morphologies; thus, the model trained on the training dataset had the same inherent bias. Our workflow could overcome these problems, and thus, the model trained on the dataset can lead to a more accurate characterisation of tumour tissues.

We further showed that the annotation accuracy increased with the number of annotated cells. In most cell types, the accuracy did not saturate, even with a large number of annotations in SegPath. One possible reason for this is that cell morphology is more diverse than what is currently known, and our dataset covers full diversity. Another possibility is that the segmentation models had a receptive field that exceeded the range of cells. Therefore, it considered the surrounding environment to make a comprehensive judgement. Therefore, it is important to create datasets with various tissues and specimens, which was an advantage of our approach for using immunostaining TMAs.

Other experimental methods exist to identify multiple cell types simultaneously in a tissue section, such as highly multiplexed IF^23^, imaging mass spectrometry^24^, and spatial transcriptomics^25^. These methods are more accurate than our approach and can identify cells that cannot be detected in H&E-stained sections. However, these methods are costly, laborious, and require additional equipment and experiments, including the optimisation of experimental conditions^26^. Segmentation from H&E-stained tissues is a complementary approach to these methods as it does not require additional equipment. More importantly, H&E staining accounts for approximately 80% of all human tissue staining performed worldwide^10^ and this method can be applied to a large number of specimens accumulated thus far. Additionally, a high-throughput analysis that will allow simultaneous comparison of multiple samples can be achieved by applying TMAs to glass slides containing fragments of tissues from numerous patients. These advantages enable comprehensive ‘pathomics’, which can be used to analyse the correlation between cell or tissue distribution and clinical information such as genomics data^27^.

The limitation of this study was that the dataset included various errors and inconsistencies derived from uneven IF staining, nonspecific staining, and errors in the cell recognition model. However, according to a previous study^28^, the supervised segmentation method is sensitive to biased errors and robust to unbiased errors. The dataset generated in our workflow is less biased than those generated by pathologists in terms of morphology. Our results further showed that the model trained on our dataset outperformed the assessments made by pathologists of several cell types, suggesting that the model can detect cells with atypical morphologies. Additionally, emerging techniques for robust learning under random label noise, such as constrained reweighting, can be used to develop more accurate segmentation models^29, 30^. We will make this large-scale dataset publicly available to enhance pathology-based cancer research and segmentation algorithm development. We plan to expand the datasets to include more cell types and lead to finer segmentation. Our approach will enhance high-throughput computational pathology by adding more information, such as the tissue context, rather than the image level category, and could eventually lead to improved diagnostic techniques and drug development for cancer patients.

## Acknowledgements

We thank Shin Aoki for helping us stain tissue specimens. We thank Editage (https://www.editage.jp/) for the English language review. This study was supported by the AMED Practical Research for Innovative Cancer Control under Grant Number JP 21ck0106640 to S.I., the AMED Project Focused on Developing Key Technology for Discovering and Manufacturing Drugs for Next-Generation Treatment and Diagnosis under Grant Number JP 21ae0101074 to S.I., and JSPS KAKENHI Grant-in-Aid for Scientific Research (B) under Grant Number 21H03836 to D.K.

## Author Contributions

Conceptualization, S.I. and D.K.; Methodology, D.K, K.S. and H.O.; Software, D.K. and T.O.; Validation, D.K., T.O.; Formal Analysis, D.K.; Investigation, D.K., T.O. and S.I.; Resources, T.O., T.U.; Data Curation, D.K., T.O., M.O. and H.K.; Resources, K.S., T.I., and T.U.; Annotation, M.H., M.O, and R.H.; Writing - Original Draft D.K., T.O., and S.I.; Writing - Review & Editing, D.K. and S.I.; Visualization, D.K.; Supervision, S.I.

## METHODS

### Sample preparation and image data acquisition

All histopathological specimens used in the generation of SegPath were obtained from patients who were diagnosed between 1955 and 2018 and had undergone surgery at the University of Tokyo Hospital. TMAs for various cancers (including glioma, meningioma, ependymoma, kidney renal clear cell carcinoma, lung adenocarcinoma, lung squamous cell carcinoma, breast adenocarcinoma, gastric adenocarcinoma, colon adenocarcinoma, pancreatic adenocarcinoma, cholangiocarcinoma, hepatocellular carcinoma, oesophageal squamous cell carcinoma, head and neck squamous cell carcinoma, urothelial tumours, bladder cancer, prostate adenocarcinoma, sarcoma, melanoma, uterine cancer, ovarian tumours, and testicular germ cell tumours) were constructed from the formalin-fixed, paraffin-embedded (FFPE) tissue blocks used for pathological diagnoses. The TMA FFPE blocks were cut to obtain 3-μm-thick sections. All histopathological specimens were anonymised in an unlinkable manner; thus, the requirement for informed consent was waived. This study was approved by the Institutional Review Board of the University of Tokyo. Information on the histopathological specimens is summarised in Supplementary Table 1.

To create the SegPath dataset, we obtained histopathological images of both H&E- and IF-stained sections from the same TMAs as follows. For H&E staining, the sections were de-paraffinised and rehydrated by immersion in xylene (#241-00091, FUJIFILM Wako Pure Chemical Corporation, Osaka, Japan) and ethanol (#057-00451, FUJIFILM Wako Pure Chemical Corporation), respectively. Haematoxylin (#6187-4P, Sakura Finetek Japan, Tokyo, Japan) and eosin (#8660, Sakura Finetek Japan) solutions were used for H&E staining following the manufacturer’s protocols. The stained sections were dehydrated by immersion in ethanol, followed by xylene. Glass coverslips (Matsunami Glass Ind. Ltd., Osaka, Japan) with Marinol (#4197193, Muto Pure Chemicals, Tokyo, Japan) were used to cover the stained sections. H&E staining, using the same protocol, was also performed to create WSIs for evaluating multi-cell type segmentation among resected specimen sections. WSIs of the H&E-stained sections were captured using a Hamamatsu Nanozoomer 2.0 HT whole slide scanner (Hamamatsu Photonics K.K., Shizuoka, Japan) at 40× (0.220818 µm/pixel) resolution. Next, we used the same sections of H&E-stained TMA sections for IF. The glass coverslips were removed by immersing the slides in xylene, rehydrating with ethanol, and washing with distilled water. For the destaining of H&E and antigen retrieval, the slides were autoclaved for 5 min at 120 °C and immersed in citrate buffer (pH 6.0; Abcam, Cambridge, UK). Endogenous peroxidase activity was measured using 0.3% hydrogen peroxide (Sigma-Aldrich, St. Louis, MO, USA) in methanol (#137-01823, FUJIFILM Wako Pure Chemical Corporation) for 15 min, followed by washing with distilled water. Nonspecific protein–protein reactions were blocked by incubating the sections in Antibody Diluent/Block (#ARD1001EA, PerkinElmer, Waltham, MA, USA) for 15 min at room temperature. The following primary antibodies were used, as summarised in Table 1: monoclonal mouse IgG anti-pan-cytokeratin, clone AE1/AE3 (without dilution; IS05330-2J; DAKO, Carpinteria, CA, USA); monoclonal mouse IgG anti-human αSMA, clone 1A4 (1:200 dilution; M085129-2; DAKO); monoclonal mouse IgG anti-human CD45RB, leucocyte common antigen, clones 2B11 + PD7/26 (1:200 dilution; IR75161-2J; DAKO); monoclonal mouse IgG anti-human N-terminal ERG, clone 9FY (without dilution; PM421AA; Biocare Medical, Concord, CA, USA), monoclonal mouse IgG anti-glycophorin A, clone JC159 (1:200 dilution; MA5-12484; Thermo Fisher Scientific, Waltham, MA, USA), polyclonal rabbit anti-human myeloid cell nuclear differentiation antigen (1:1000 dilution; HPA034532-100UL; Sigma-Aldrich); monoclonal rabbit IgG anti-human MIST1/bHLHa15 protein, clone D7N4B (1:100 dilution; #14896; Cell Signaling Technology, Beverly, MA, USA); polyclonal mouse anti-human CD3 (1:200 dilution; IS50330-2J; DAKO); and monoclonal mouse IgG anti-human CD20cy, clone L26 (1:200 dilution; IS60430-2J; DAKO). IF staining using each of the aforementioned primary antibodies (AE1/AE3, αSMA, CD45, ERG, glycophorin A, MNDA, MIST1, and CD3/CD20 mix) was performed for 2 h at 4 °C, according to the instructions of Opal Multiplex IHC KIT (#NEL811001KT; PerkinElmer). The Opal Polymer HRP solution (ARH1001EA; PerkinElmer) was used to enhance the signals by incubating the sections for 10 min at room temperature. The sections were then incubated with 100 µL of Opal 690 fluorophore (1:10 dilution; FP1497001KT; PerkinElmer) at room temperature for 10 min to achieve 690-nm single-color IF staining. Nuclear staining was performed with DAPI solution (FP1490A; PerkinElmer) at room temperature for 5 min. The ss were then covered with glass coverslips (Matsunami Glass Ind. Ltd.) using Prolong Gold antifade reagent with DAPI (P36931, Thermo Fisher Scientific). WSIs of the IF staining TMA slides were captured using a Hamamatsu Nanozoomer 2.0 HT whole slide scanner at 40× (0.220818 µm/pixel) resolution.

### WSI pre-processing

Large artefacts (i.e., tissue folds and air bubbles) in each WSI were marked by pathologists before analysis. In the patch extraction process, patches overlapping the marked regions or heavily blurred regions with a variance of the Laplacian filter^31^ < 0.0005 in the grayscale image were removed. Additionally, tissue region candidates were extracted from grayscale H&E slides at zoom level 4 (1/16 of the 40× resolution) by applying Otsu binarisation after Gaussian blur with an 81 × 81-kernel. Connected regions within the size of 12.8–256 million pixel^2^ at 40× resolution were regarded as the tissue regions. Patches within the tissue regions of 1,024 × 1,024 pixels with a stride of 1,024 pixels were then extracted. The patches within 200 pixels at 40× resolution from the edges of the tissue regions were discarded as nonspecific IF staining is often observed at the edge of the tissue^32^.

### Image registration and patch extraction

To create masks for the deep learning model of H&E-stained histological images, each IF image was registered to the H&E-stained image of the same slide. Image registration was performed using a multistep procedure that began with coarse WSI level registration and proceeded to fined-grained, patch-level registration. Nuclear regions were considered in the calculation to accurately align the two images. Specifically, the haematoxylin colour component extracted using the scikit-image’s ‘rgb2hed’ function in the H&E image and DAPI channel component in the IF image were used for registration. First, discrete Fourier transform (DFT)-based rigid registration was performed to estimate the optimal vertical and horizontal translation between H&E WSI and paired IF WSI at zoom level 6 (1/64 of the 40× resolution). After the patch pairs of 1024 × 1024 pixels at zoom level 1 (1/2 of the 40× resolution) had been extracted from the same position of the aligned WSI pairs, the DFT-based rigid registration was performed again to obtain a finer-grained registration, and the vertical and horizontal translation levels were recorded. Kernel density estimation using Gaussian kernels was applied to the two-dimensional distribution of the translations, and the vertical and horizontal translation levels with the highest densities were used to register all image pairs in the same WSI. Subsequently, 1024 × 1024-pixel tiles at 40× resolution were extracted again from the aligned WSI pairs. After two additional rounds of the same DFT-based rigid registrations at zoom levels 1 and 0 at 40× resolution, nonrigid registration using the Demons algorithm^33^ was applied after the histogram matching filter. We used a multiresolution pyramid with three layers (with shrinkage factors of 8, 4, and 2 and a smoothing sigma of 12, 8, and 4). A gradient descent with a learning rate of 1.0 and 20 iterations was used for parameter optimisation. Finally, 20-pixel margins from the edges were removed, such that the image did not include unregistered regions.

### Initial mask generation

For mask generation, it is necessary to determine the cut-off values for positive IF signals and remove false-positive signals due to artefacts, registration errors, or nonspecific signals from blood cells.

Inconsistencies between the intensities of the DAPI nuclear channel in the IF image and the haematoxylin component in the H&E-stained image, indicating the existence of artefacts or registration errors, were detected by calculating the Pearson’s correlation coefficient between the two signal intensities. Patches with correlation coefficients smaller than 0.5 were removed for further analysis. False-positive signals derived from the autofluorescence of RBCs were removed by masking the positively predicted regions using the RBC segmentation neural network trained on the anti-CD235a antibody stained dataset. Based on visual inspection, an IF signal intensity > 50 (epithelium, smooth muscle, and RBC) or 25 (others) was regarded as a positive signal in the initial mask generation step.

For the epithelia, smooth muscle, and RBCs, the positive signal area was used as a segmentation mask without modification. For leucocytes, myeloid cells, lymphocytes, plasma cells, and endothelial cells, positive signals from the target cells were transferred into the nuclei based on the IF staining pattern to obtain a more consistent result and improve the interpretability of the segmentation model. Cellpose^12^ version 0.6.5 was applied to the DAPI nuclear channel in the IF images to detect the nuclei. We selected a model with the following parameters: diameter = 30, channels = [3,0], batch_size = 64, and cellprob_threshold = 0.1. Nuclei were masked if over 40% of them contained positive signals. Finally, one iteration of morphological erosion with a 3 × 3-kernel was applied to each region of the nuclei to prevent multiple cells from sticking together, which could cause an underestimation of the cell count.

For deep neural network training, all patches were divided into training, validation, or test sets so that all patches from the same TMA spot belonged to the same set. TMA spots in each TMA were detected as clusters by applying the DBSCAN clustering algorithm^34^ implemented in scikit-learn to patches using the x and y coordinates as the input features, maximum distance set to 3,000 pixels, and min_samples set to 5. The validation and test sets contained patches from two TMA spots in each TMA slide, and the rest were placed into the training set.

### Training deep neural network for segmentation

The encoder–decoder neural networks were trained for semantic segmentation. The combination of the encoder and decoder was independently optimised for each cell type or tissue. The backbone of the encoder was a pretrained convolutional neural network, such as ResNet^35^ trained on the 2012 ILSVRC ImageNet dataset, or EfficientNet^36^ trained on 300 million unlabelled images from JFT^37^ using noisy student training^38^. The decoder module was selected from one of three models: U-net^39^, U-net++^40^, or DeepLabV3+^41^. The network was trained using randomly sampled patches with sizes of 984 × 984 pixels and batch sizes of 16. During training, the weights in all layers of the decoders and segmentation head were updated through the RAdam optimiser with a weight decay of 1 × 10^-4^, *β*_1_ = 0.9, and *β*_2_ = 0.999. Data augmentation and normalisation were applied in the following order:

- Random crop to 640 × 640 pixels

- Colour, contrast, and brightness augmentation (hue [-0.1, 0.1], saturation [0.9, 1.1], contrast [0.9, 1.1], and brightness [0.9, 1.1])

The data were normalised to mean = [124.0, 116.0, 104.0] and std = [58.6, 57.3, 57.6]

- Random horizontal and vertical flips

- Random affine transformation with rotation with up to 180 degrees, and scale with scaling factor ranging from 0.9 to 1.1 with reflection padding

-Random Gaussian blur of a 3 × 3-kernel with probability = 0.3

The backbone and architecture of the deep learning model and hyperparameters, including the learning rate, were optimised using the tree-structured Parzen estimator algorithm^42^ based on the validation Dice score. The validation Dice score was evaluated across all images at once instead of averaging the Dice scores of each patch as the positively stained areas varied drastically among patches. The hyperparameters optimised in this study are listed in Supplementary Table 2. All segmentation models were trained for 25 epochs. At least five trials were tested, and the model with the best validation Dice score was chosen for the subsequent analysis.

### Improvement of cut-off intensity and nucleus overlap ratio

The cut-off values of the signal intensities (=25 or 50) and nucleus overlap rate (40%) in the initial mask generation may not be optimal. Since we observed heterogeneity in the signal intensities of some of the sections, a single cut-off value for the signal intensity across one TMA slide may not be appropriate. Otsu’s binarisation is often applied in similar scenarios, but it is not easy to differentiate patches with overall low signals because of weak staining of positive cells or the absence of positive cells, and the latter results in many false-positive masks.

We observed that the staining strength gradually changed in the section. The segmentation network could detect positive cells with a certain level of accuracy, even if it was trained on the initial mask. Based on this observation, we iteratively improved the cut-off values. First, linear regression analysis was carried out to detect patches with positive cells by setting the intercept to zero, where the explanatory value was the IF intensity. The dependent variable was the cell probability of the trained deep neural network model, both of which were smoothed by Gaussian blur with a 3 × 3- kernel. Patches with a regression coefficient > 1 and maximum IF intensity > 10 without RBC regions were considered positive. For each positive patch, the initial cut-off values were determined by applying Otsu’s binarisation to the patch and the nearest eight positive patches. To avoid extreme cut-off values, they were clipped to a minimum of 10 and a maximum of 50. For the epithelia, the cut-off was reduced by 20% as we observed the heterogeneous staining of anti-pan-CK antibodies between the cytoplasm and nuclei, with weaker signal intensities in the nucleus. Finally, the thresholds for each patch, including the negative patch, were determined using the weighted average cut-off values of the nearest 16 positive patches. A Gaussian distance weight of 1/3000 pixel from the target patch was used for the weight. For leucocytes, myeloid cells, lymphocytes, plasma cells, and endothelial cells, the nucleus overlap cut-off, which maximises Matthew’s correlation coefficient between the prediction and mask, was used. In contrast to the signal intensity cut-off, the same cut-off value was adopted for each cell type. Based on the new segmentation masks, the segmentation networks were trained again, and the cut-off intensity and nucleus overlap ratio were optimised. These processes were repeated twice, and we verified that the mask remained almost unchanged after the second optimisation.

### Annotation by pathologists

For each cell type, except RBCs, 10 patches were randomly selected from the training data. Three trained pathologists independently performed the annotation task for the patches using the Labelbox annotation tool (San Francisco, CA, USA)^43^. Tissue regions were selected with polygonal annotations, whereas cells were selected with point annotations to the centre of the nuclei. In the first round, only H&E images were shown to the pathologists and annotated. Regions or cells selected by at least two pathologists were used as the HE-path for subsequent experiments. In the next round, both H&E and IF images without DAPI overlaid with H&E images were shown to the same pathologists and annotated again. Regions or cells selected by at least two pathologists were used as pGT data. For point annotations to the cells, annotations by two pathologists were regarded as overlapping if they were within an 8-pixel distance (≒1.77 µm).

#### Evaluation of masks and predictions

The accuracy of the annotations was evaluated based on the Dice coefficient between the pGT and HE-path, IF-mask, or prediction. The pixel- and cell-level Dice coefficients were calculated for tissue and cell segmentations, respectively.

The HE-path and IF-mask or prediction were regarded as overlapping if any point in the HE-path was on the IF-mask or predicted region.

### Morphological evaluations

We evaluated the morphological parameters of the annotated cells: cartwheel-shaped nuclei for plasma cells, nucleus density for lymphocytes, and distance from the nearest RBC to the endothelial cells. The presence of typical cartwheel-shaped nuclei in each plasma cell was determined by a pathologist. For lymphocyte intensity, the nucleus regions were annotated by a pathologist using Labelbox, and the mean intensity of the haematoxylin component estimated by the rgb2hed function in the scikit-image was used for the evaluation. For the distance from the nearest RBC to the endothelial cells, all RBCs were annotated by a pathologist using Labelbox, and the pixel distance between each endothelial cell and nearest RBC was used for the evaluation.

### Effect of training data size on the segmentation performance

Patches were individually sampled in the training set until the number of patches or cells reached the predetermined value, as shown in Supplementary Table 3. This process was repeated thrice for each value, except for the entire training set. Using these datasets, the deep neural network models were trained and tested on the same test set.

We trained a U-net with a resnet18 backbone pretrained on ImageNet on the training dataset using the same procedure for all evaluation datasets. The network was trained using randomly sampled patches with a size of 960 × 960 pixels and batch size of 16. During training, the weights in all layers of the decoders and the segmentation head were updated using the RAdam optimiser with a learning rate of 0.01, weight decay of 1 × 10^-4^, *β*_1_ = 0.9, and *β*_2_ = 0.999. The same augmentation and normalisation were applied as described in the previous section. The Dice loss was used as the loss function. The model was trained for a maximum of 10,000 epochs. If the validation Dice coefficient did not increase for five consecutive epochs, early stopping was applied, and the best model based on the validation Dice coefficient was used for testing.

### Validation cohort

WSIs from two institutes were used for the validation of multi-cell-type segmentations of H&E-stained images. Specimens were obtained from 1) three patients with gastric adenocarcinoma who underwent surgery and were diagnosed at the University of Tokyo Hospital between 1955 and 2018 and 2) four patients with salivary gland tumours (salivary duct carcinoma, Warthin’s tumour, and cystadenoma) who underwent surgery and were diagnosed at Tokyo Medical and Dental University Hospital between 1990 and 2020. Resected specimens of gastric adenocarcinoma were prepared from the FFPE blocks and sliced to a thickness of 6 μm. All histopathological specimens were anonymised. This study was approved by the Institutional Review Board of each university.

### Multi-cell-type segmentation

Based on the deep neural network model for the segmentation of each tissue or cell type, we performed multi-cell type segmentations of the H&E-stained images. To consider the lineage hierarchy of cells or tissues, the segmentation results were merged, beginning with coarse categories then overwritten with fine-grained categories. The four groups were defined as follows and overwritten in the below order. The following layer and label encoding were adopted:

[layer 0] 0: background; 1: stroma (other than smooth muscle cells).

[layer 1] 2: epithelial cells; 3: smooth muscle cells.

[layer 2] 4: leucocytes, 5: endothelial cells, 6: red blood cells.

[layer 3] 7: lymphocytes, 8: plasma cells, 9: myeloid cells.

*x_ij_*, 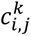, and 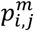 denote the pixel intensity after grayscale conversion, predicted label at the *k*-th layer, and output logit value of the segmentation model for *m*-th cell type at the (*i,j*)-th pixel in the image, respectively.

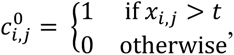

where *t* is Otsu’s threshold for the WSI based on the pixel intensity after grayscale conversion. The predicted label was updated using the following recursive calculation:

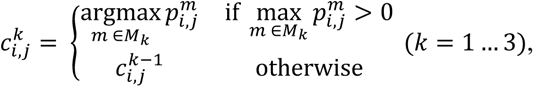

where *t* is the set of cell types in the *k*-th layer, and 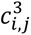 was used as the prediction label.

We applied the model ensemble approach for each tissue or cell type using 1 to 3 different models with the best validation Dice coefficients during training. To obtain the WSI level segmentation results, inference was executed for a 7680 × 7680-pixel patch from the WSI, and the result of each patch was assembled into a WSI.

### Implementation details

Rigid and nonrigid image registrations were performed using imreg version 2.0.1a (https://github.com/matejak/imreg_dft) and SimpleITK version 2.0.2 Python library^44^, respectively. Kernel density estimation was performed using SciPy version 1.3.1^45^. The neural networks were trained using Python version 3.8.5, PyTorch Lightning version 1.4.2 (https://www.pytorchlightning.ai/), and Segmentation Models Pytorch version 0.2.0 (https://github.com/qubvel/segmentation_models.pytorch). To speed up the model training, mixed precision (16-bit) training implemented in PyTorch Lightning was used. Hyperparameter optimisation was performed using the Optuna version 2.7.0^46^. The training and testing were performed on the NVIDIA DGX-1 server with 8 NVIDIA Tesla V100 GPUs and 256 GB RAM.

---

**Extended Data Fig. 1.**
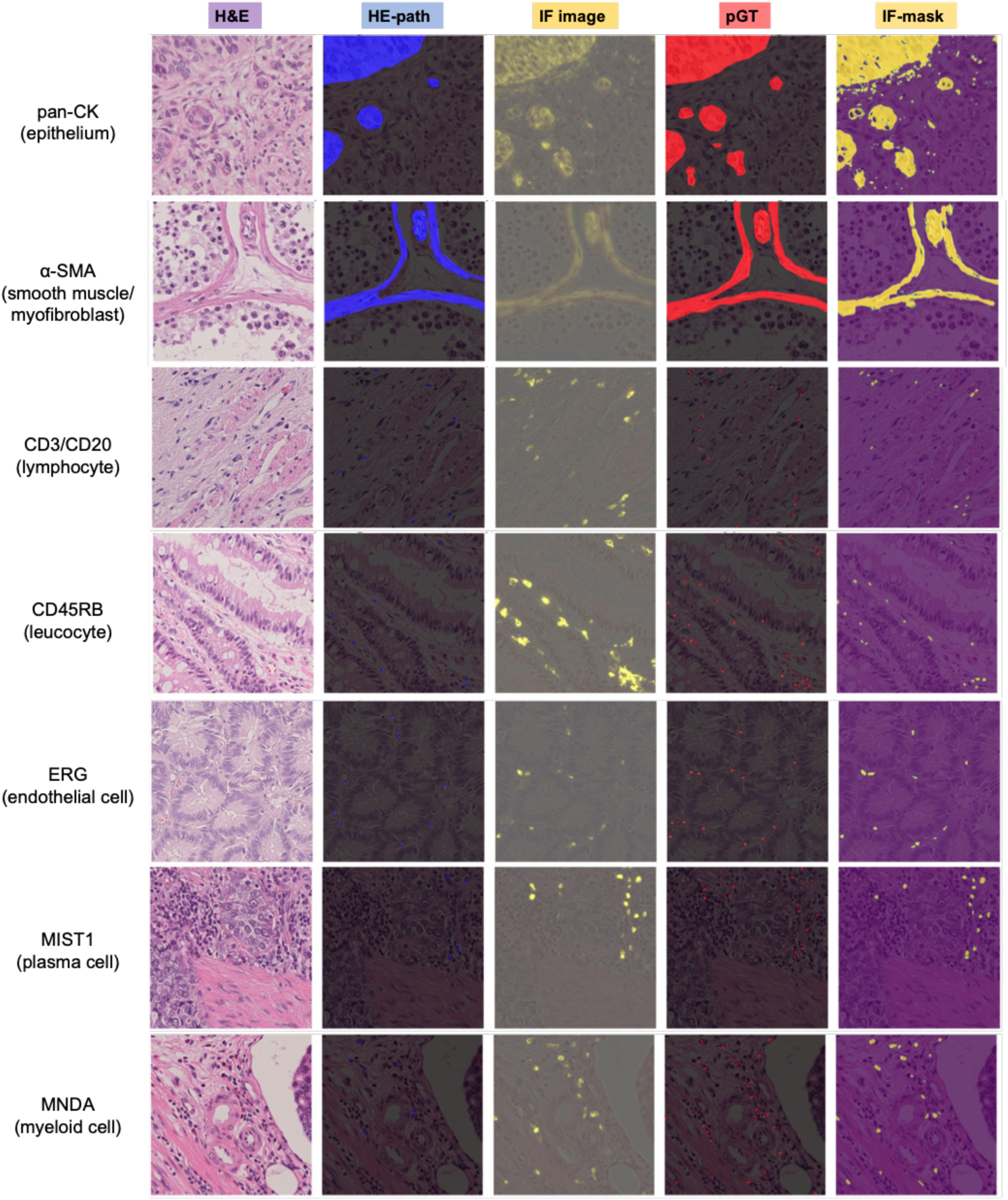
Annotation samples. H&E images (first column) and the corresponding annotations for pathologist annotations based on the HE-path images (second column), IF images (third column), HE-path and IF images (fourth column), and IF-mask (fifth column). From the second to fifth columns, the images were overlaid with the corresponding H&E images.

**Extended Data Fig. 2.**
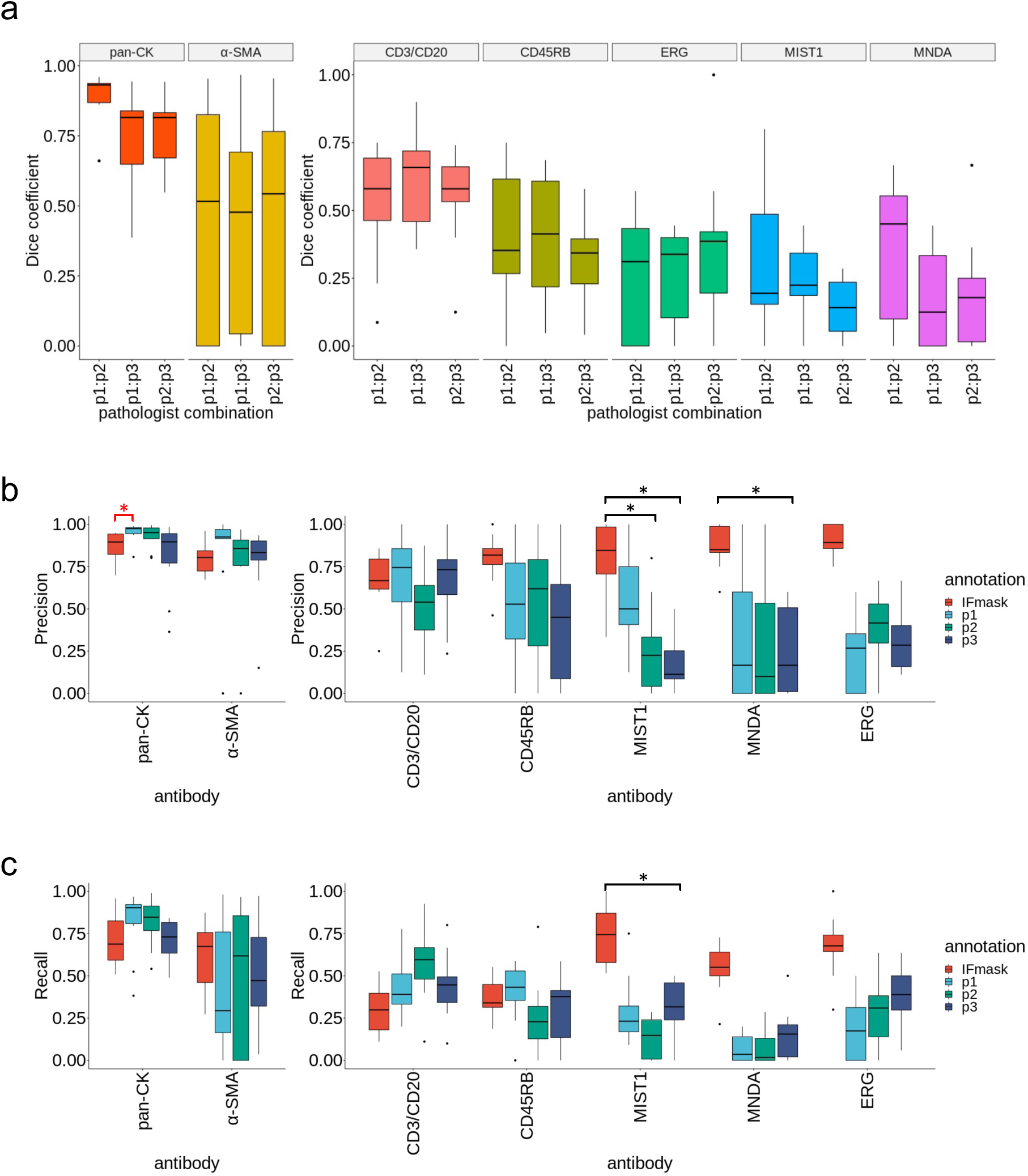
Evaluation of the annotation accuracy of SegPath. **a,** Inter-pathologist concordance of the annotations in terms of Dice coefficients (*n* = 10 patches of 217.5 × 217.5 µm for each tissue or cell type). Three pathologists annotated the tissues or cells based on the H&E images only. **b, c,** Comparison of the annotation accuracies between pathologists and the automatically generated masks in terms of precision shown in **b** and recall shown in **c**. (*n* = 10 patches of 217.5 × 217.5 µm for each tissue or cell type). pGT annotations were performed by pathologists who examined both the H&E and corresponding IF images. Regions or cells annotated by at least 2 of 3 pathologists were used. \**P* < 0.05. The red asterisk indicates that the precision of Pathologist 1 was significantly higher than that of the IF-mask.

**Extended Data Fig. 3.**
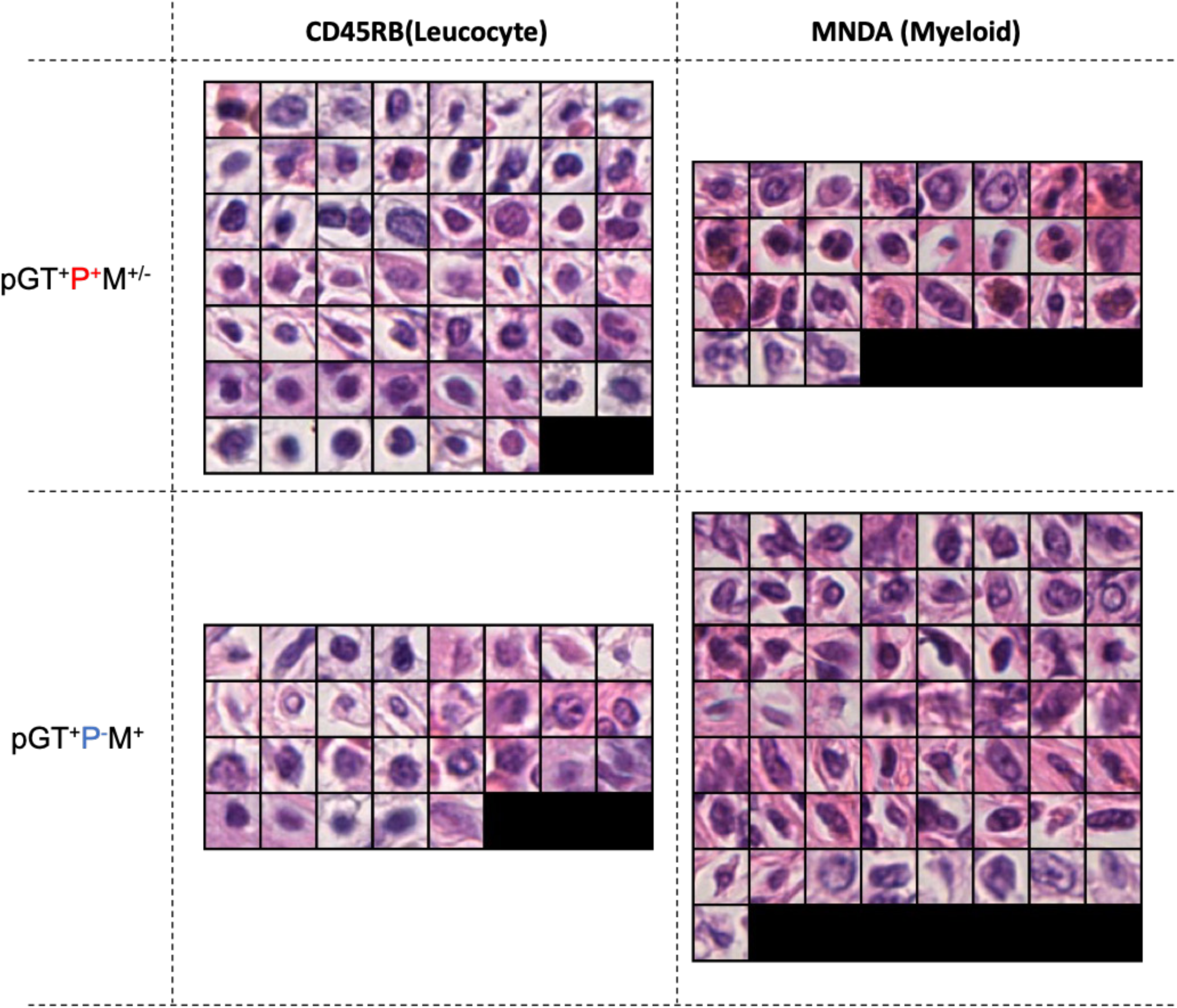
Annotated cell images for leucocytes and myeloid cells. pGT cell images annotated by multiple pathologists (pGT^+^P^+^M^+-^) and not identified by multiple pathologists but successfully annotated by the masks (pGT^+^P^-^M^+^). pGT, ground truth; P, HE-path; M, IF-mask.

**Extended Data Fig. 4.**
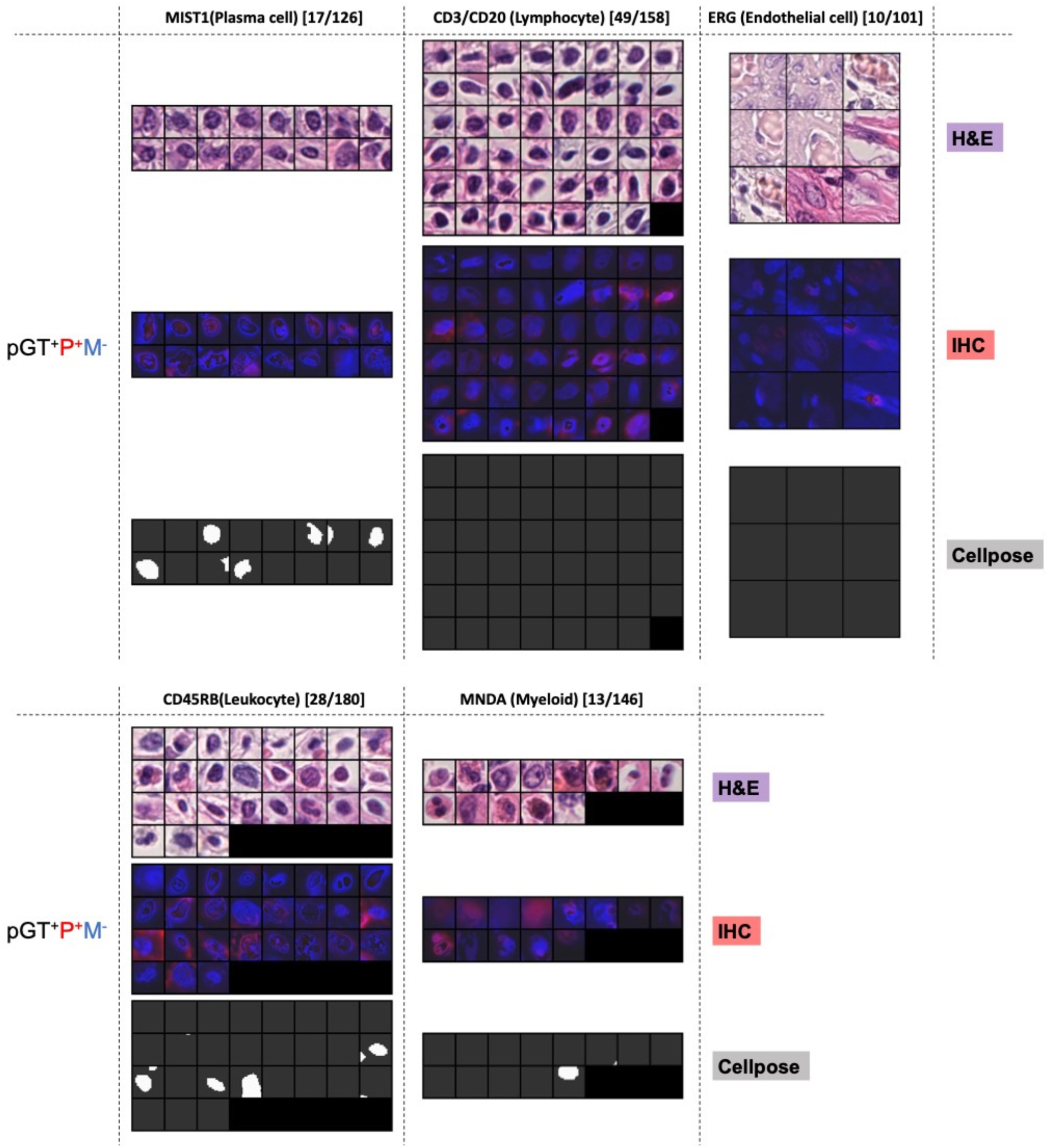
Error analysis of cells correctly identified by pathologists but missed by IF-masks. pGT cell images were annotated by multiple pathologists but were not identified by the masks (pGT^+^P^-^M^-^). The corresponding IF images of the cell-specific antibodies (red), DAPI nuclear staining (blue), and nuclei detected by Cellpose (white) are shown. The numbers in brackets indicate the number of cells correctly identified by pathologists, but missed by the IF-masks, and the number of pGT cells. pGT, ground truth; P, HE-path; M, IF-mask.

**Extended Data Fig. 5.**
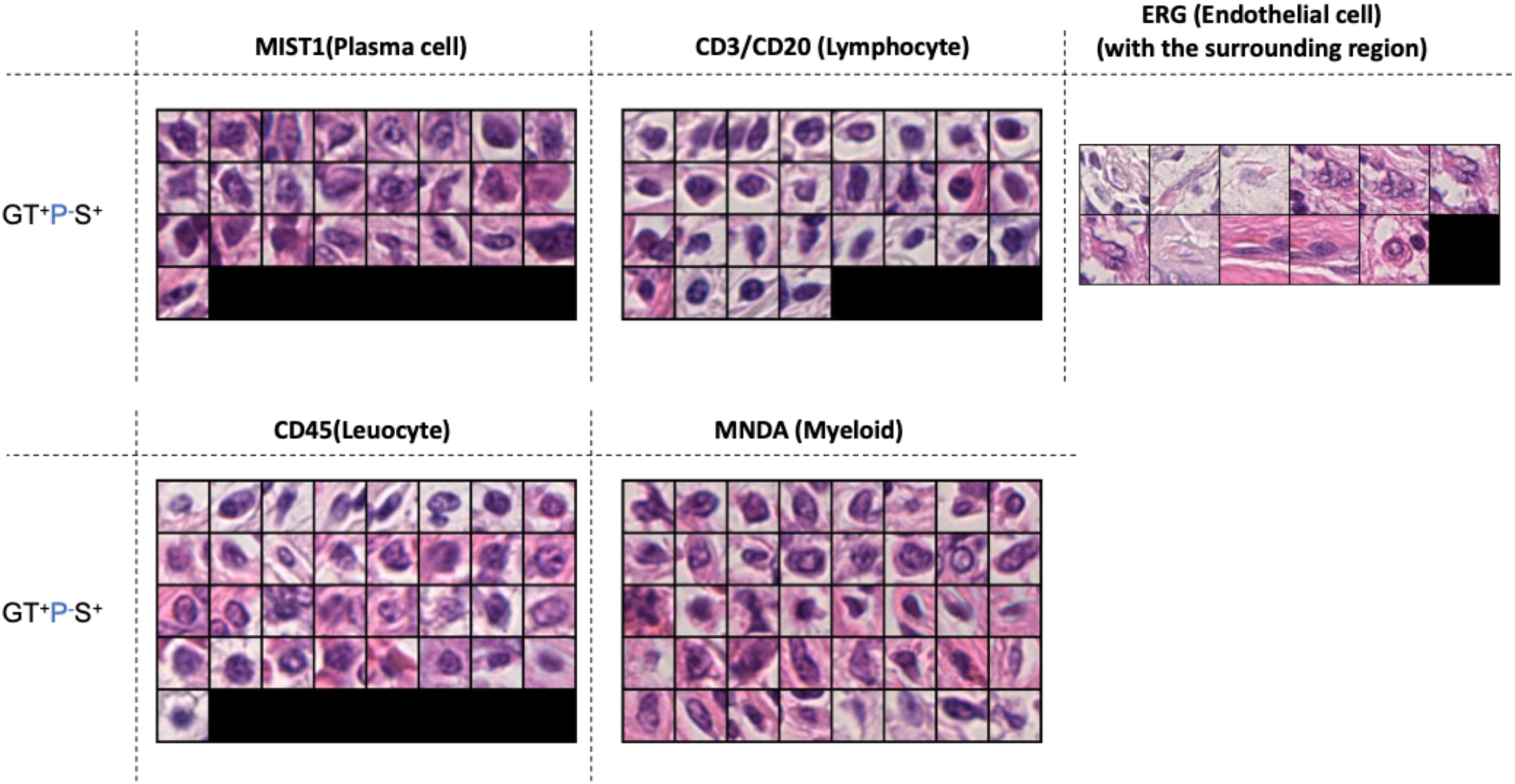
Annotated cell images. pGT cell images not annotated by multiple pathologists but successfully predicted by the trained segmentation models (top, pGT^+^P^-^S^+^), and those identified by multiple pathologists but not predicted by the segmentation models (pGT^+^P^+^S^-^). pGT, ground truth; P, HE-path; S, Prediction using the segmentation model.

**Extended Data Fig. 6.**
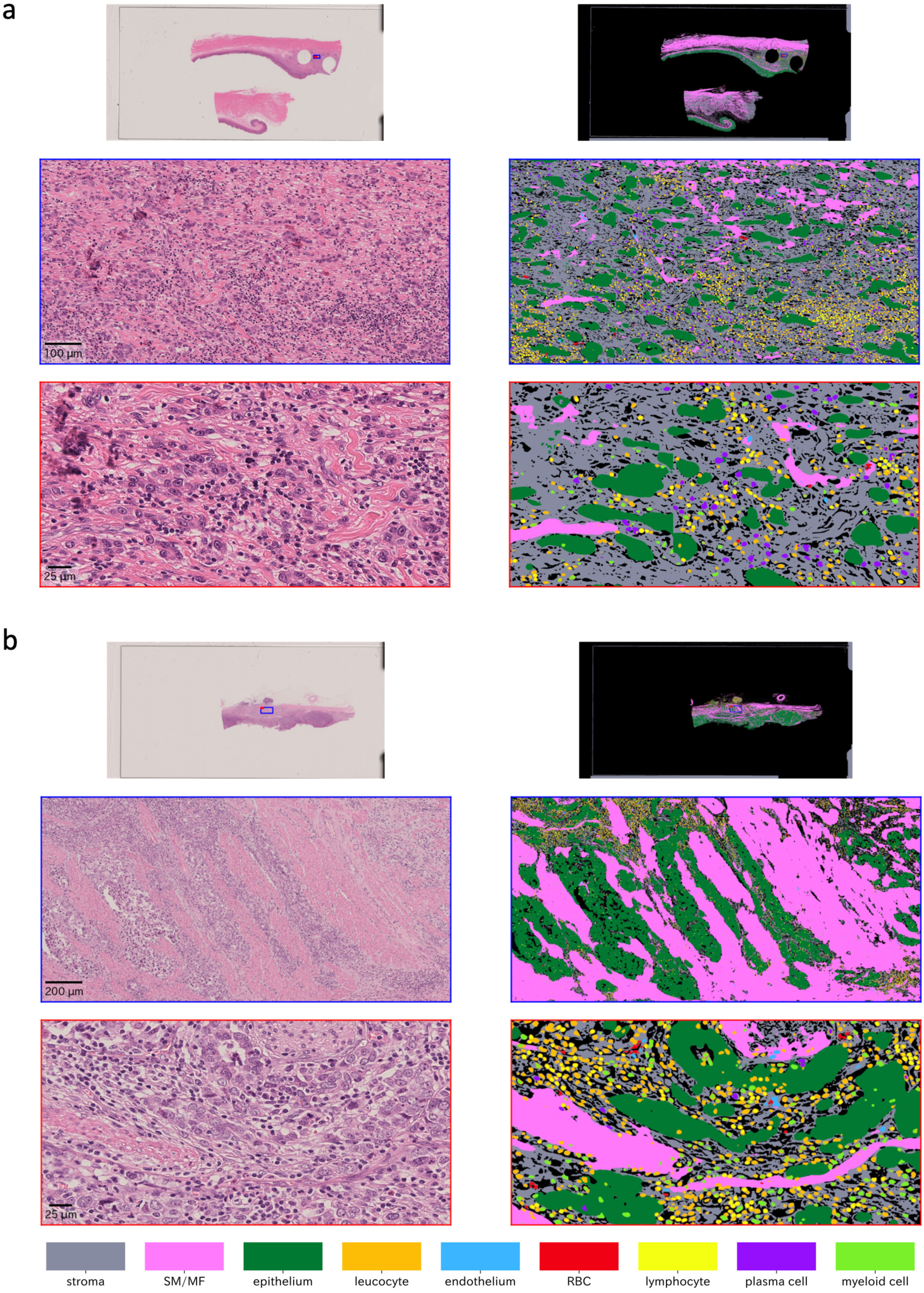
Visualisation of the multi-cell/tissue segmentation results by the trained segmentation models (continued from. Fig. 6e**). a, b,** Gastric cancer cases. SM, smooth muscle; MF, myofibroblast. The regions in the blue and red rectangles in the top WSI level image are enlarged in the middle and bottom images with a bounding box of the same colour.

**Extended Data Fig. 7.**
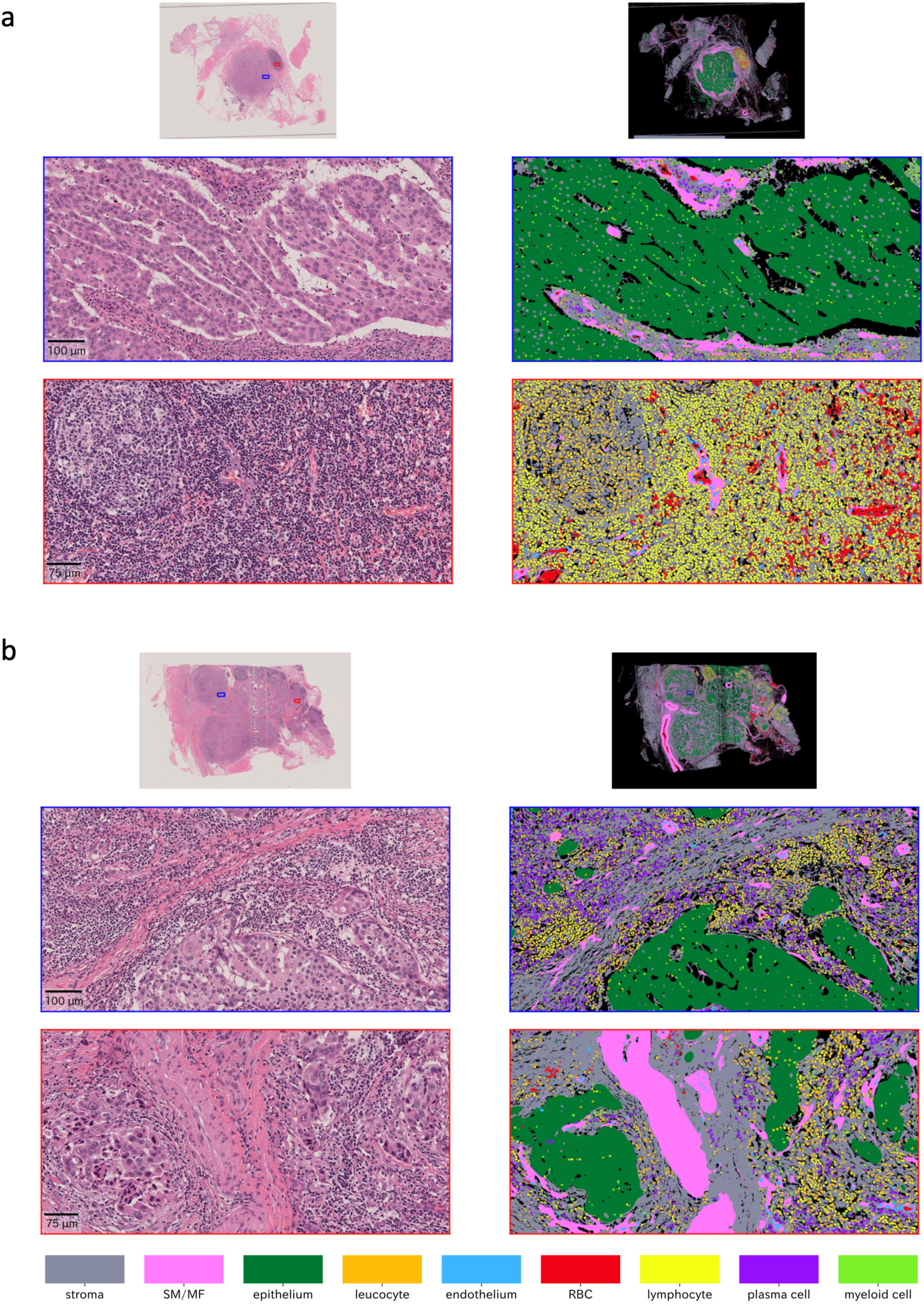
Visualisation of the multi-cell/tissue segmentation results by the trained segmentation models (continued from. Fig. 6e). Malignant salivary gland tumours. **a, b,** Salivary duct carcinoma cases. SM, smooth

**Extended Data Fig. 8.**
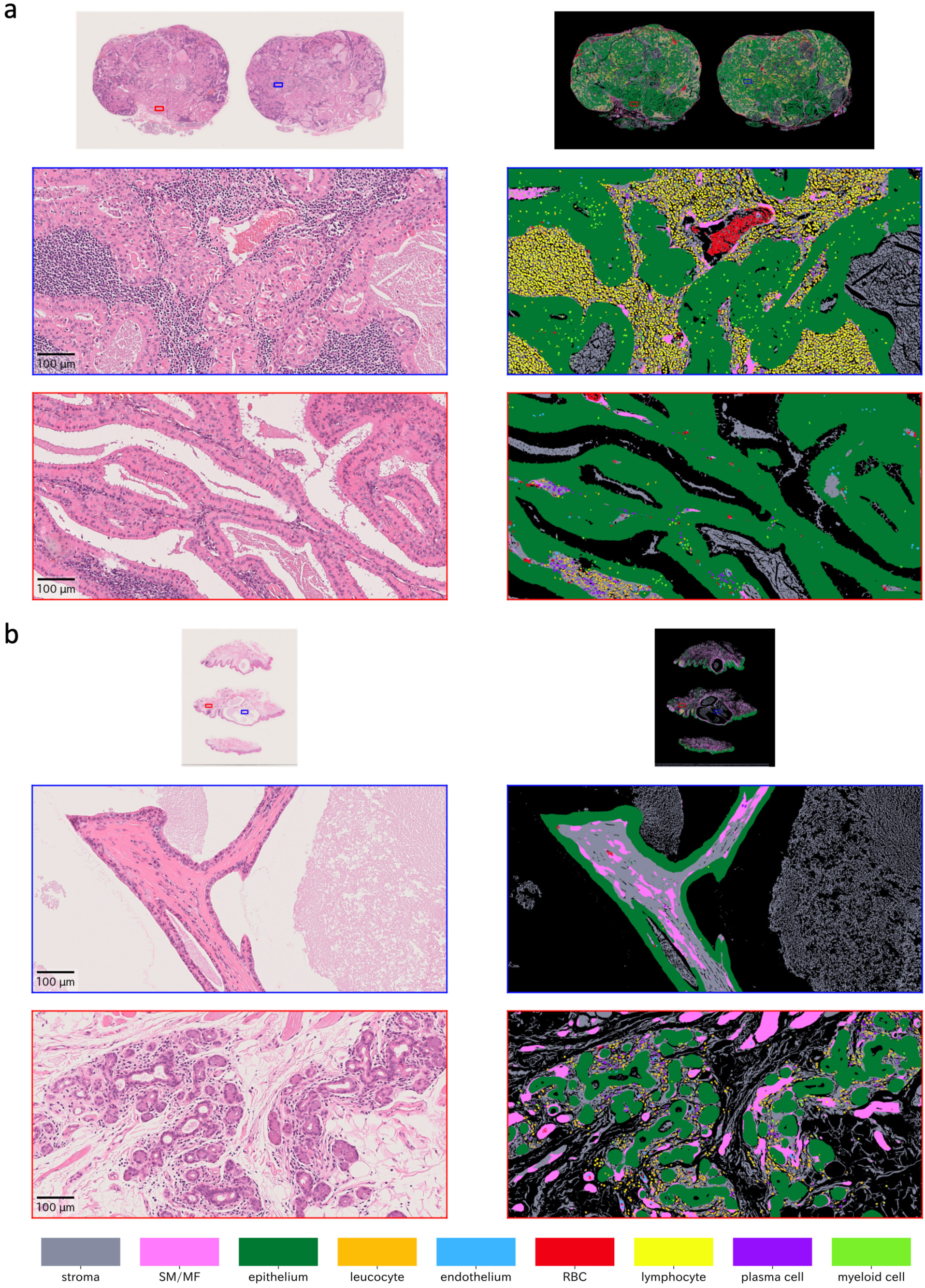
Visualisation of the multi-cell/tissue segmentation results by the trained segmentation models (continued from. Fig. 6e). Benign salivary gland tumours. **a,** Warthin’s tumour. **b,** Cystadenoma. SM, smooth

