## Supplementary Figures for "Beyond pathologist-level annotation of large-scale cancer histology for semantic segmentation using immunofluorescence restaining"

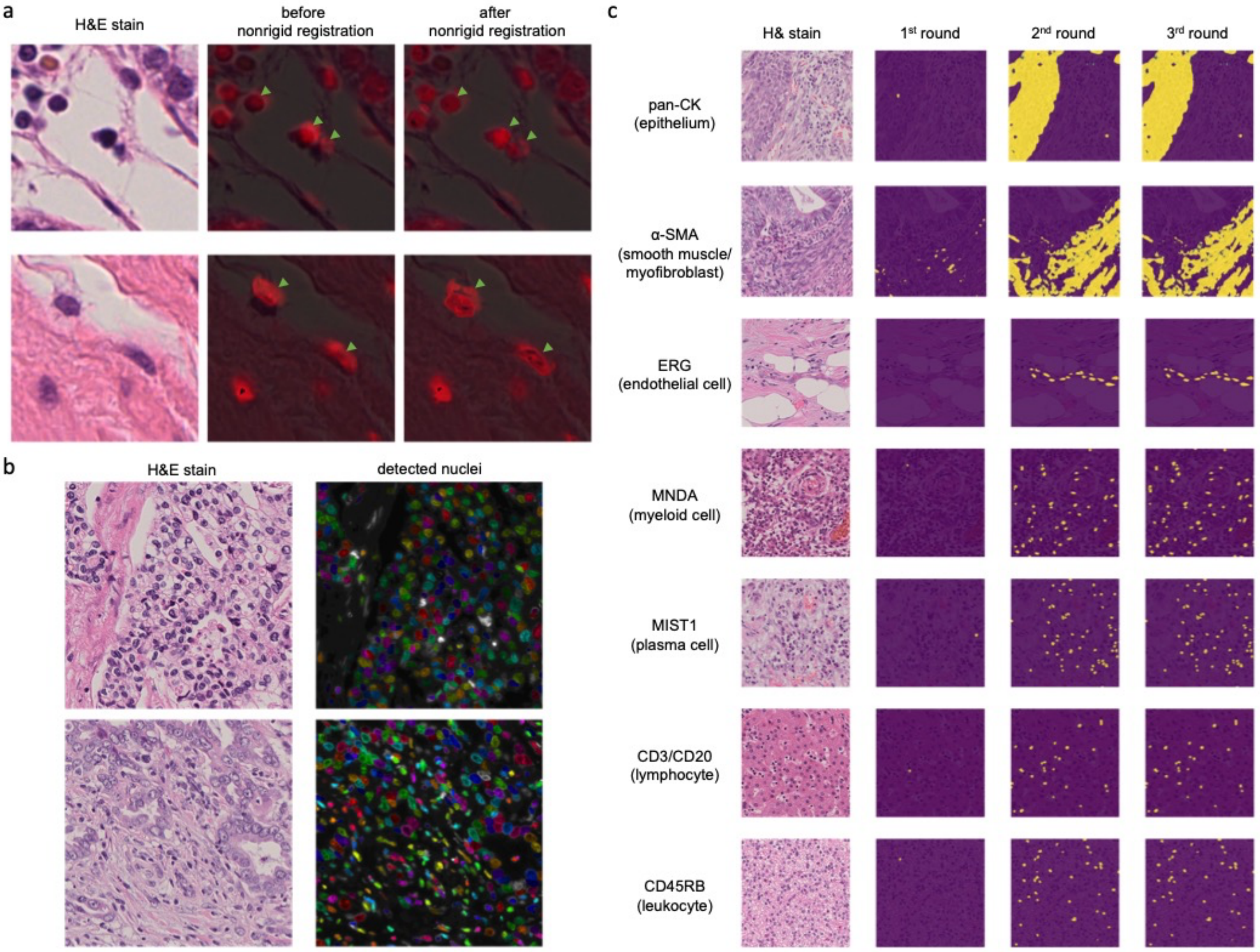


**Supplementary Fig. 1. Pre-processing of H&E and IF sections. a,** H&E-stained and DAPI channel images before and after nonrigid registration between estimated haematoxylin components in H&E. H&E images are overlaid in the DAPI channel images. The DAPI component is shown in red, which was converted from blue to improve visibility, in the middle and right images. Arrowheads indicate cells drastically corrected by nonrigid registration. **b,** H&E-stained image and detected nuclei in DAPI components in the IF-stained sections. Coloured regions represent detected nuclei, and grey regions represent nucleus candidates that did not reach the detection threshold. **c,** IF threshold improvement in each phase.

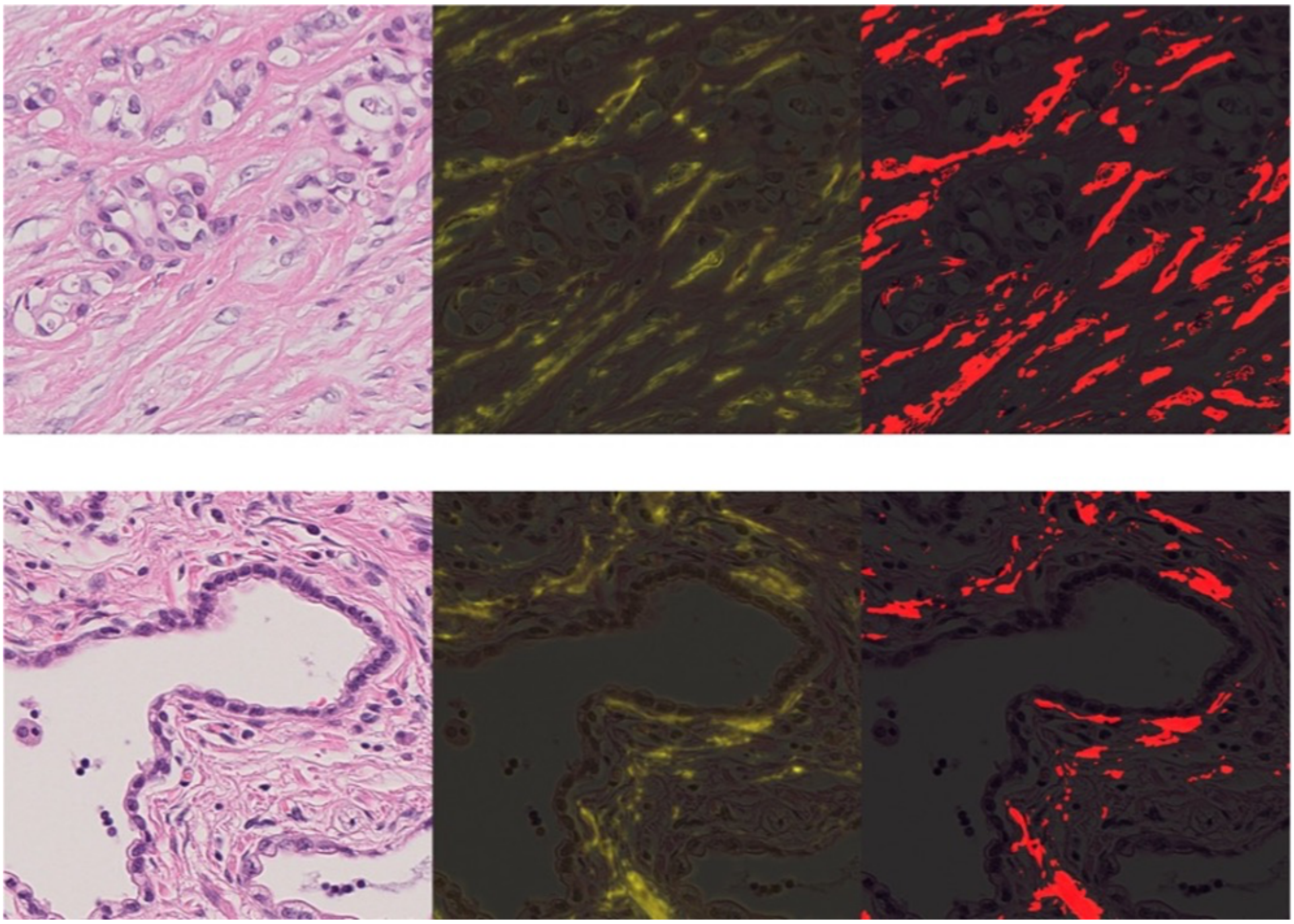

**Supplementary Fig. 2. Weak staining of αSMA.** Each triplet shows an H&E-stained image, the corresponding registered IF image, and generated mask image (positive regions are indicated in red), from left to right. The organs are shown above each triplet. All image patches are 217.5 × 217.5 µm.

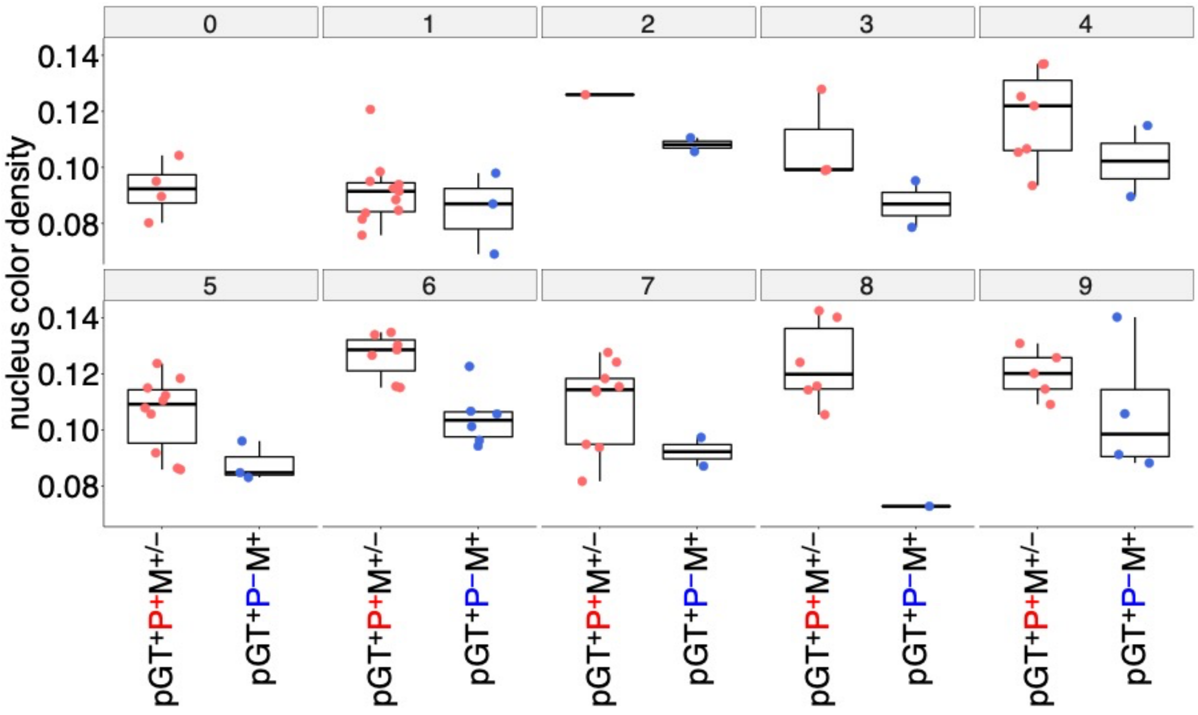


**Supplementary Fig. 3. Nuclear haematoxylin intensities of lymphocytes for each patch.** pGT, ground truth; P, HE-path; M, IF-mask.

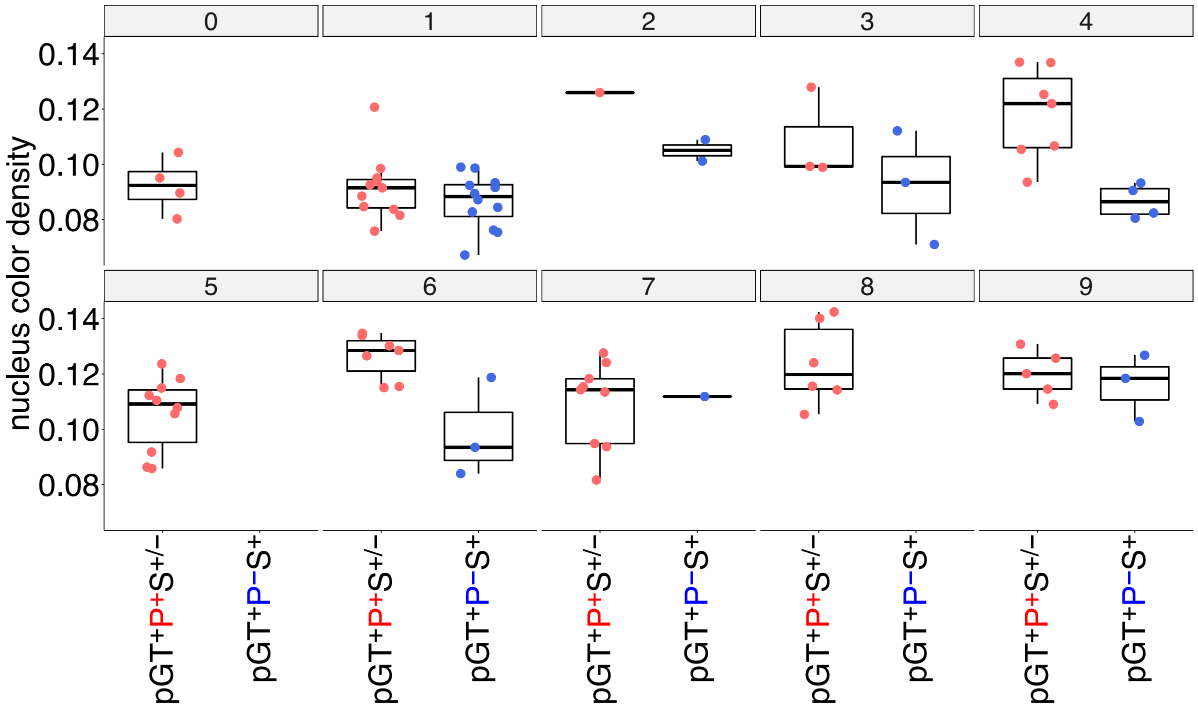


**Supplementary Fig. 4. Nuclear haematoxylin intensities of lymphocytes.** pGT, ground truth; P, HE-path; S, Prediction by the segmentation model.

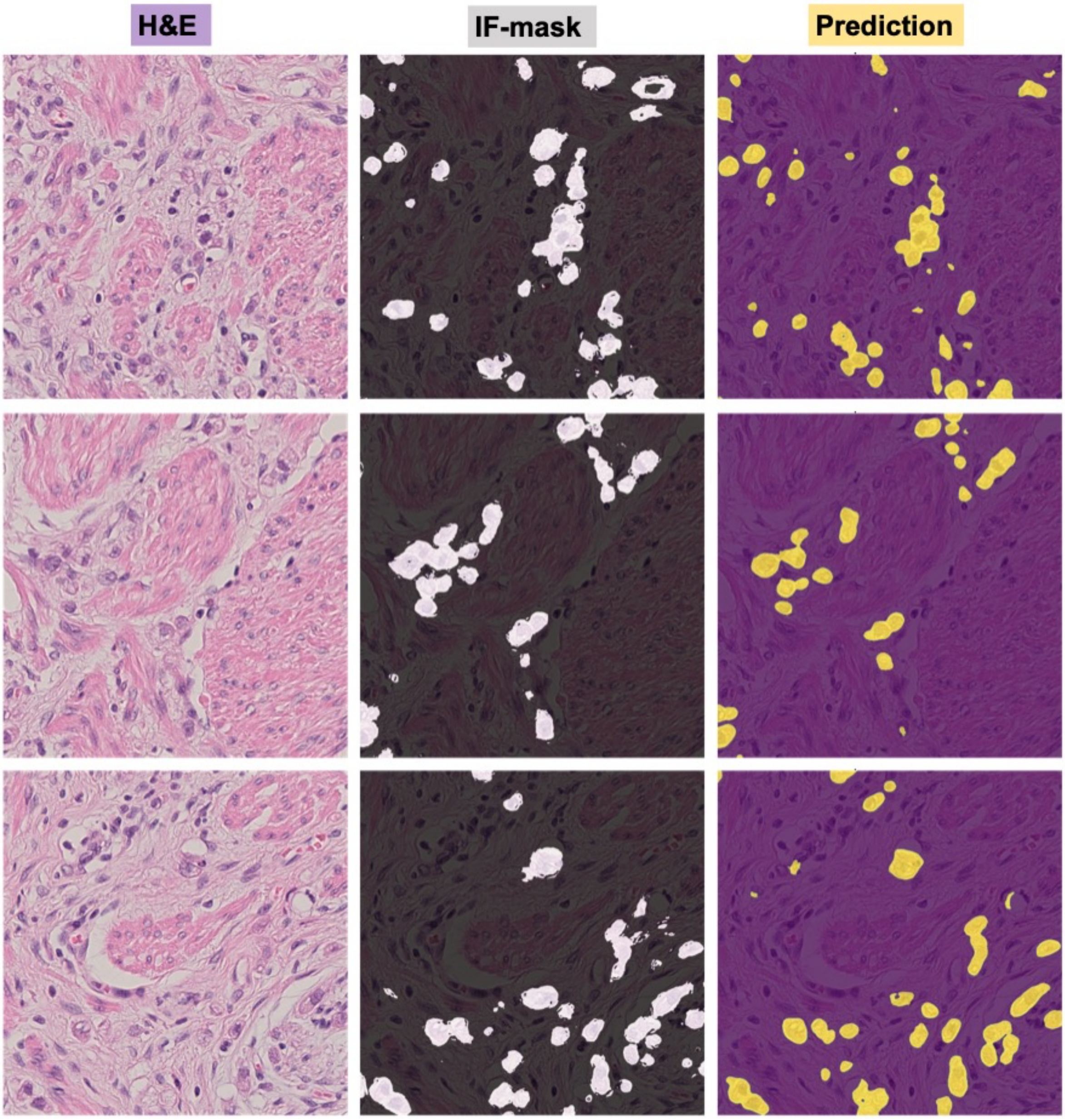


**Supplementary Fig. 5. Isolated gastric cancer cells.** H&E-stained images (first column), IF-masks (second column), and predicted epithelium regions by the trained segmentation model (third column) shown for three different regions.
